# Obesity-Associated Changes in Immune Cell Dynamics During Alphavirus Infection Revealed by Single Cell Transcriptomic Analysis

**DOI:** 10.1101/2024.10.10.617696

**Authors:** Muddassar Hameed, Andrea R. Daamen, Md Shakhawat Hossain, Sheryl Coutermarsh-Ott, Peter E. Lipsky, James Weger-Lucarelli

## Abstract

Obesity induces diverse changes in host immunity, resulting in worse disease outcomes following infection with various pathogens, including arthritogenic alphaviruses. However, the impact of obesity on the functional landscape of immune cells during arthritogenic alphavirus infection remains unexplored. Here, we used single-cell RNA sequencing (scRNA-seq) to dissect the blood and tissue immune responses to Mayaro virus (MAYV) infection in lean and obese mice. Footpad injection of MAYV caused significant shifts in immune cell populations and induced robust expression of interferon response and proinflammatory cytokine genes and related pathways in both blood and tissue. In MAYV-infected lean mice, analysis of the local tissue response revealed a unique macrophage subset with high expression of IFN response genes that was not found in obese mice. This was associated with less severe inflammation in lean mice. These results provide evidence for a unique macrophage population that may contribute to the superior capacity of lean mice to control arthritogenic alphavirus infection.

**Graphical abstract:** 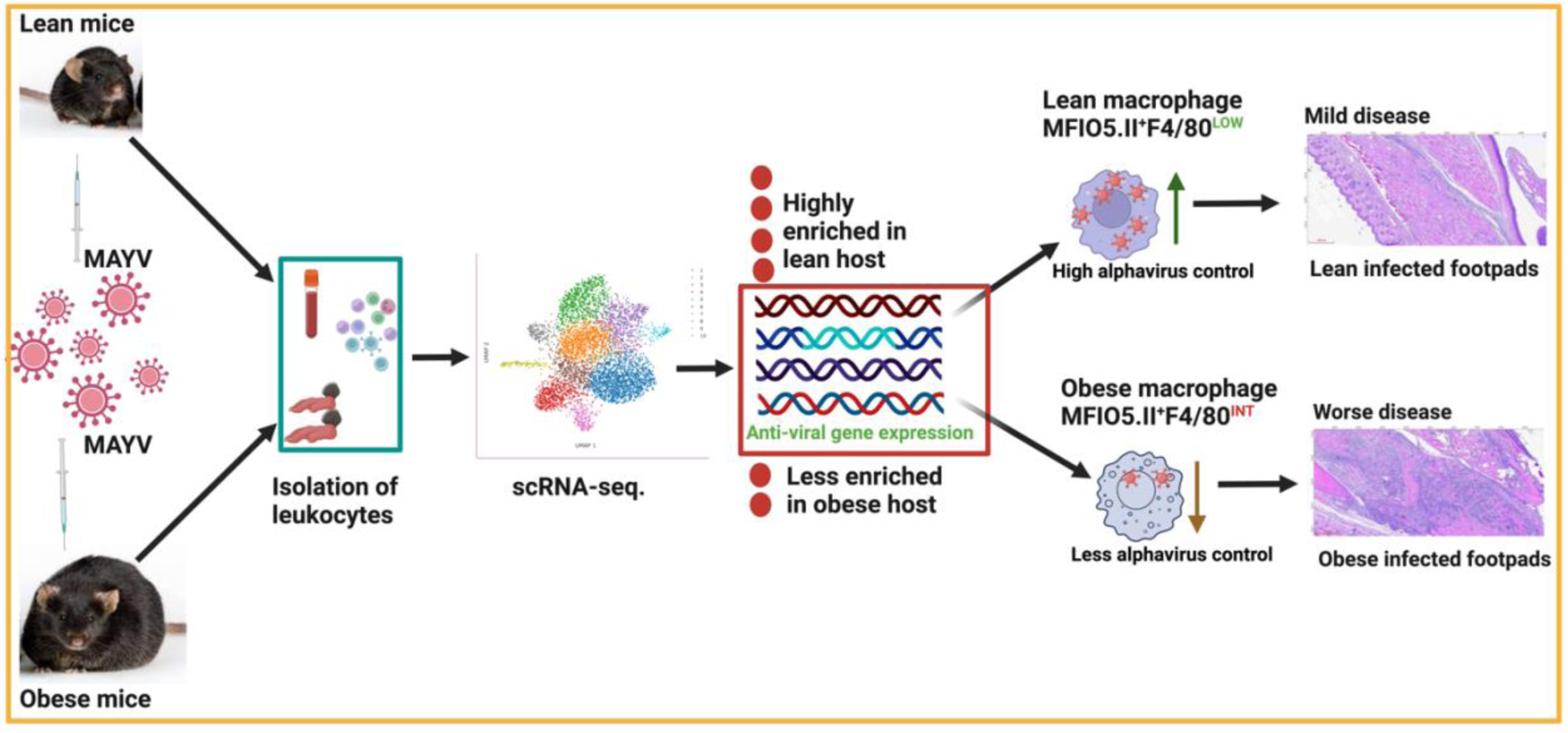

**Highlights:**

- Obesity worsens disease outcomes following arthritogenic alphavirus infection.
- Arthritogenic alphavirus infection causes significant shifts in immune cell populations in the blood and footpad.
- Blood monocytes from lean mice had higher expression of interferon response genes at the later stage of infection.
- Footpads in lean mice have an expanded population of F4/80lo macrophages with an intense interferon response gene signature before and after alphavirus infection that is not found in obese mice.
- Macrophages in obese mice express lower levels of interferon response genes, have a unique necroptosis signature, and higher F4/80 expression.

## Introduction

Arthritogenic alphaviruses, including chikungunya virus (CHIKV), Mayaro virus (MAYV), and Ross River virus (RRV), cause acute and chronic disease in humans^1^. Acute disease is characterized by fever, rash, myalgia, and arthralgia^2,3^. Roughly 40% of infected people develop chronic disease associated with debilitating symptoms of chronic arthritis/arthralgia and fatigue that can last for years after infection^3–5^. A recent study suggested that mortality is higher than previously estimated following CHIKV infection, with an average case-fatality rate of 0.8 deaths per 1,000 cases in Brazil^6^. Despite their impact on human health, no approved therapeutics are available to treat alphavirus-induced disease. Although the FDA recently approved a vaccine to prevent chikungunya disease, its real-world effectiveness remains to be documented^7^.

Epidemiological reports have identified factors associated with severe chikungunya disease in humans, including obesity^8^, which is expected to affect 50% of the United States population by 2030^9^ and currently impacts 13% of adults worldwide^10^. Consistent with this, we previously showed that obese mice develop more severe disease following infection with CHIKV, MAYV, or RRV^11^; however, the mechanisms underlying the difference in disease outcomes remain unexplored. Obese hosts develop a chronic, low-grade inflammation related to the ongoing release of pro-inflammatory cytokines such as tumor necrosis factor (TNF) and interleukin 6 (IL-6)^12–14^. This results in a dysfunctional immune response to various infectious diseases^15–17^, making obese hosts more susceptible to severe disease, including during infection with arthritogenic alphaviruses^11^.

Arthritogenic alphavirus disease has been associated with a dysregulated immune response, with various cell types and pro-inflammatory cytokines implicated in pathogenesis^18,19^. Inoculation of mice with an arthritogenic alphavirus induces footpad swelling, myositis, and bone loss, similar to the disease observed in humans^20–23^. However, there is an incomplete understanding of the immune processes driving immune/inflammatory responses to these viruses. Previous efforts to understand disease pathogenesis in arthritogenic alphavirus-infected mice have included transcriptomic analysis, largely based on bulk RNA sequencing^24–26^, which offers limited insights into the heterogeneity of individual cell types^27,28^. Recently, scRNA-seq has emerged as a means of assessing the contribution of individual cell types to pathogenesis^29–32^ and providing valuable insights into the contribution of individual immune cell types to disease processes.

Here, we carried out scRNA-seq from immune cells isolated from blood during peak viremia (2 days post-inoculation [dpi]) and from blood and footpads at the time of peak footpad swelling (7 dpi) from mock and MAYV-infected lean and obese mice. We identified large shifts in immune cell populations in the blood and footpads following MAYV infection. Notably, B cells dramatically decreased in the blood at 2 dpi, whereas monocytes were enriched in both lean and obese mice. In the footpad, macrophage populations increased considerably. Most cell types showed significant enrichment for genes in the type I and II interferon (IFN) pathways following MAYV infection. We identified a unique subset of macrophages (CSF1R^+^SiglecF^-^ F4/80^+lo^MHCII^+high^) that predominated in the footpad both before and following MAYV infection in lean mice; these cells were not identified in obese mice. This population in lean mice was enriched for IFN response genes. Since serum and tissue levels of IFNs were similar in lean and obese mice, the decreased expression of IFN-response genes in obese mice suggests a defect in response to IFN. These results suggest a specific cellular and functional abnormality in obese mice that is associated with more severe disease.

## Results

### Differences in clinical severity and immune cell dynamics during MAYV infection in lean and obese hosts

To profile the host immune response to arthritogenic alphavirus infection, we infected lean and obese mice with MAYV in the footpad. MAYV is a BSL2 virus and causes similar disease in humans and mice as CHIKV, which requires stricter BSL3 containment. As we previously demonstrated^11^, MAYV infection caused more weight loss in obese mice compared to lean controls (Figure 1A). MAYV infection also caused greater footpad swelling in obese mice compared to lean mice (Figure 1B), which was associated with pronounced inflammatory cell infiltration and skeletal muscle and soft tissue inflammation (Figure 1C). Finally, we observed significantly greater inflammation and composite footpad pathology scores for obese mice compared to the lean group (Figure 1D). These results confirm that obese mice develop more severe disease following MAYV infection than lean mice.

**Figure 1.**
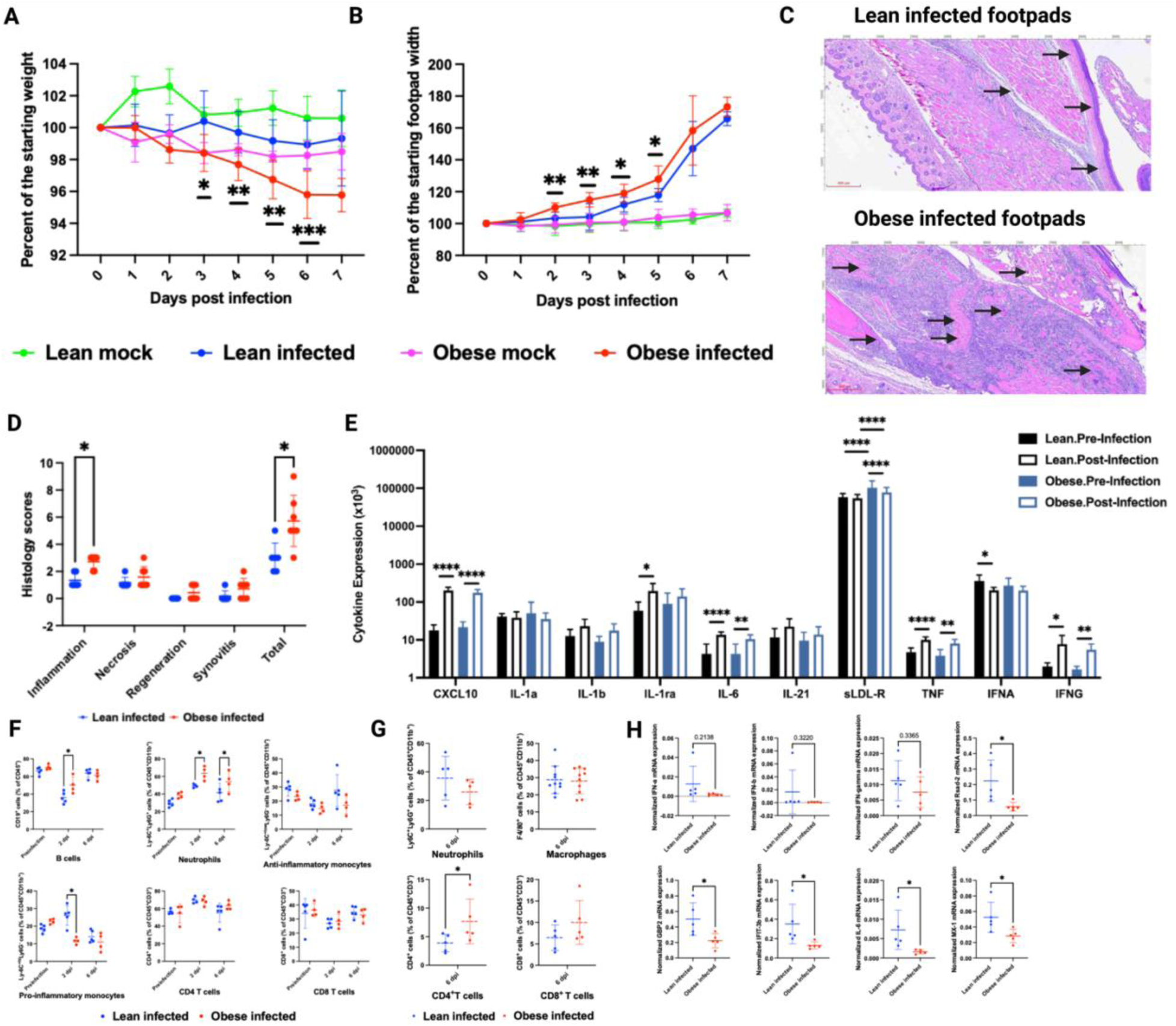
Clinical severity and immune cell profiling following arthritogenic alphavirus infection in lean and obese mice. C57BL/6J mice were fed a 10% fat (lean) or 60% fat (obese) diet for 18-20 weeks. Mice were infected with 10^4^ PFU of MAYV strain TRVL 4675 through injection of both hind footpads. (**A-B**) Weight and footpad swelling were measured daily. Data are presented as the percent of baseline body weight (**A**) and footpad width (**B**) (two experiments, n = 9-10). Data were analyzed by two-way ANOVA with Tukey’s correction compared to the lean infected group. The error bars represent the standard deviation, bars indicate mean values, and asterisks indicate statistical difference; *p<0.05, **p<0.005, ***p<0.001. (**C-D**) At seven days post-infection, the right hind footpad was collected and fixed. The fixed tissues were then sectioned and stained with hematoxylin and eosin (two experiments, n=10) (**C**); areas with pronounced infiltrates are indicated by arrows. Images were captured at 20X; scale bar=400 μm. (**D**) An independent anatomic pathologist scored the tissues in a blinded manner. Data were analyzed using multiple unpaired t-tests with the Holm-Sidak correction for multiple comparisons. The error bars represent the standard deviation, bars indicate mean values, and asterisks indicate statistical difference; *p<0.05. (**E**) Multiplex Luminex cytokine assay comparison between pre-infection and 2 days post infection in lean and obese mice. *p<0.05, **p<0.01, ***p<0.001, ****p<0.0001. See Figure S1 for the expression of other cytokines. (**F**) Dot graphs present the frequency of B cells (CD45^+^CD19^+^), neutrophils (CD45^+^CD11b^+^Ly6G^+^Ly6C^+^), anti-inflammatory monocytes (CD45^+^CD11b^+^Ly6G^-^Ly6C^+low^), pro-inflammatory monocytes (CD45^+^CD11b^+^Ly6G^-^Ly6C^+high^), CD4^+^ T cells (CD45^+^CD3^+^CD4^+^), and CD8^+^ T cells (CD45^+^CD3^+^CD4^+^) at pre-infection, 2, and 6 dpi in the blood of lean and obese MAYV-infected animals detected by flow cytometry. Data were analyzed using multiple unpaired t-tests with the Holm-Sidak correction for multiple comparisons. The error bars represent the standard deviation, bars indicate mean values, and asterisks indicate statistical difference, (one experiment, n = 4-5), *p<0.05. (**G**) Dot graphs present the frequency of neutrophils (CD45^+^CD11b^+^Ly6G^+^Ly6C^+^), macrophages (CD45^+^CD11b^+^F4/80^+^), CD4^+^ T cells (CD45^+^CD3^+^CD4^+^), and CD8^+^ T cells (CD45^+^CD3^+^CD4^+^) at the time of peak swelling in the footpads detected by flow cytometry. Data were analyzed using unpaired t-tests with Welch’s correction. The error bars represent the standard deviation, bars indicate mean values, and asterisks indicate statistical difference, *p<0.05. (**H**) RNA was extracted from footpads of MAYV-infected lean and obese mice at 6 dpi and subjected to RT-qPCR. See Fig S2 for the expression of other genes. Data were analyzed using unpaired t-tests with Welch’s correction. The error bars represent the standard deviation, bars indicate mean values, and asterisks indicate statistical difference, **p<0.01.

We also assessed the levels of key cytokines and inflammatory markers involved in viral response in the blood of lean and obese mice pre-infection and at 2 dpi (Figure 1E, S1). Expression of CXCL10, IL-6, TNF, and IFNG was increased with infection in both lean and obese mice, whereas expression of IL-1ra was only increased in lean-infected mice, and expression of soluble LDL-R (sLDL-R) was only increased in obese infected mice (Figure 1E). Notably, only expression of sLDL-R significantly increased in infected obese mice compared to lean mice.

Next, we carried out flow cytometric analysis on blood immune cells collected at pre-infection, 2 dpi, and 6 dpi (Figure 1F). We observed a significantly higher percentage of neutrophils (P = 0.02) and pro-inflammatory monocytes (P = 0.02) pre-infection in obese mice compared to lean mice (Figure 1F). No differences were observed for other tested cell types. At 2 dpi, obese mice had a higher percentage of neutrophils (P = 0.02) and lower pro-inflammatory monocytes (P = 0.005) than lean mice (Figure 1F). B cells trended toward a higher percentage in obese compared to lean mice (P = 0.09) (Figure 1F). There were no differences in any tested cell types between lean and obese MAYV-infected groups at 6 dpi (Figure 1F). We also performed flow cytometry on footpad immune cells isolated from lean and obese MAYV-infected mice at peak footpad swelling. There was no difference observed in the proportions of neutrophils, macrophages, and CD8^+^ T cells (Figure 1G). However, a significantly higher percentage of CD4^+^ T cells was seen in obese infected mice footpads. These data suggest that obese mice have slightly altered immune cell populations following alphavirus infection.

We next performed RT-qPCR on RNA isolated from footpads from MAYV-infected lean and obese mice to evaluate interferon and interferon-stimulated gene (ISG) expression. We observed no statistical difference in IFN-α, IFN-β, and IFN-γ expression between lean and obese groups. Interestingly, ISGs such as Rsad-2, GBP-2, IL-6, IFIT-3b, and MX-1 had significantly higher expression in MAYV-infected lean compared to obese mice (Figure 1H). Moreover, there was a trend toward higher expression of IFIT-1, IFIT-2, STAT-1, IFIT-204, IRF-1, and CXCL-10 in lean infected footpads than obese footpads (Figure S2). Overall, these data suggest that lean mice have a more robust antiviral response to arthritogenic alphavirus infection despite similar levels of IFNs.

To interrogate the impact of arthritogenic alphavirus on immune cell population dynamics and functionality more thoroughly, we next performed scRNA-seq on immune cells from the blood and footpads from MAYV-infected lean and obese mice. We isolated CD45^+^ cells at two time points, representing the peak of viremia (2 dpi) and footpad swelling (7 dpi); a graphical workflow of the experiment is presented in Figure 2A. For each time point, cells from mock- and MAYV-infected mice were integrated separately from lean and obese mice, visualized as uniform manifold approximation and projection (UMAP) plots, and clustered based on conserved marker genes. As a result, we identified 9-12 cell clusters per group at 2 dpi and 11-16 cell clusters per group at 7 dpi in blood samples (Figure S3 and S4). Clusters were then annotated based on the expression of immune cell-specific genes (Figure 2B-C). As expected, a comparison of mock-infected lean and obese mice revealed that immune cell populations remained unchanged between 2 and 7 dpi in the blood (Figures 2D and 2F). In contrast, we observed a considerable shift in B cells, monocytes, neutrophils, T cells, and eosinophils between 2 and 7 dpi in blood samples of MAYV-infected lean and obese mice compared to mock-infected controls (Figure 2E and 2G). In lean MAYV-infected mice, B cells, neutrophils, and T cells increased from 2 dpi to 7 dpi, whereas monocytes and eosinophils were reduced (Figure 2D-E). Similar changes were also observed in B cells and monocytes from obese MAYV-infected mouse blood from 2 to 7 dpi (Figure 2F-G). Overall, these data highlight the considerable impact of MAYV infection on immune cell population dynamics during disease progression in lean and obese hosts.

**Figure 2.**
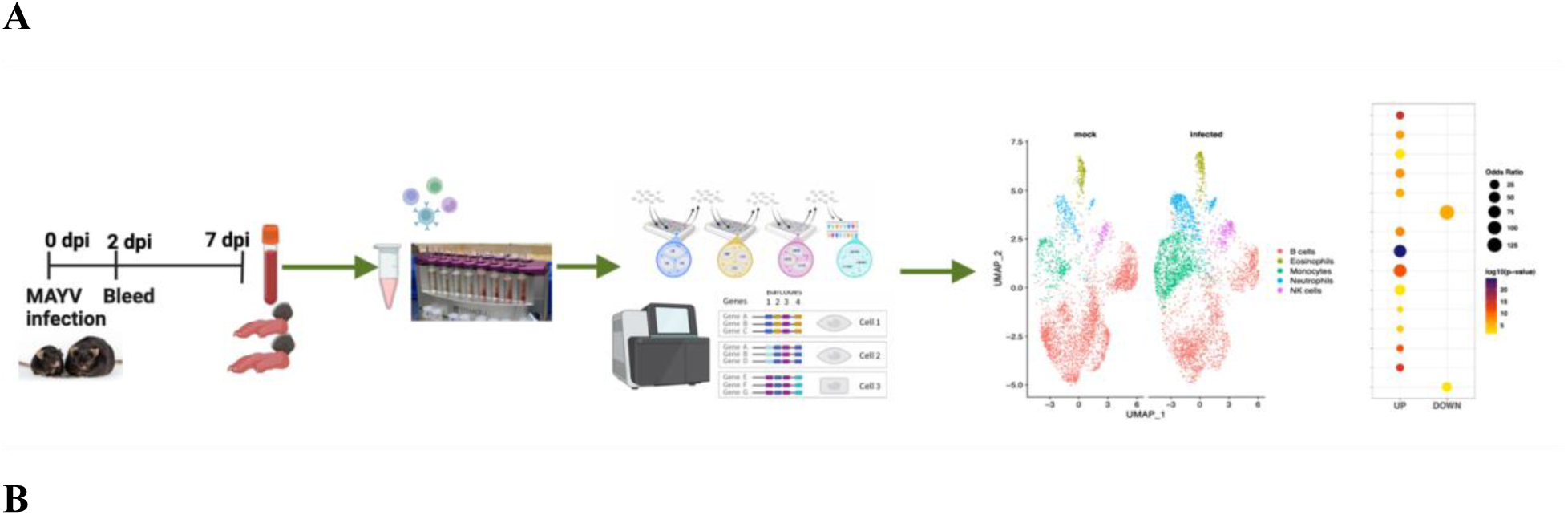
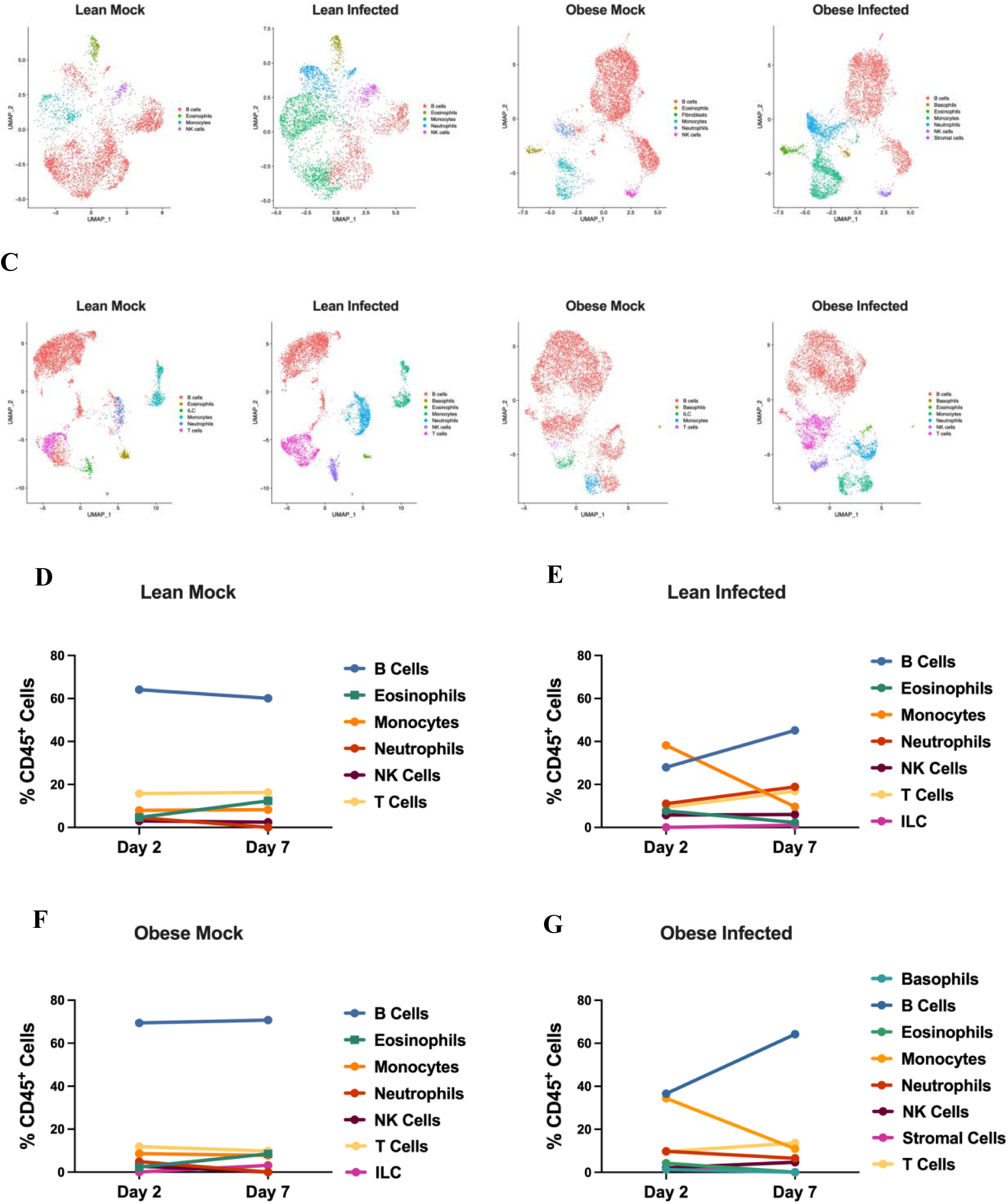
Single cell transcriptomics of immune cell dynamics following arthritogenic alphavirus infection in lean and obese mice. C57BL/6J mice were fed a 10% fat (lean) or 60% fat (obese) diet for 18-20 weeks. Mice were infected with 10^4^ PFU of MAYV strain TRVL 4675 through injection of both hind footpads. Blood immune cells were collected at 2 and 7 dpi and processed for scRNA-seq. See Figure S3 for the graphical workflow of experiment. **A. Graphical workflow of scRNA-sequencing.** Mice were inoculated with MAYV or viral diluent, and blood and footpads were collected. Then, leukocytes were isolated, and CD45^+^ cells were purified through positive selection. Libraries were prepared for scRNA-seq, which was followed by data analysis. (**B-C**) Uniform Manifold Approximation and Projection (UMAP) showing integrated datasets from blood at 2 dpi (**B**) and 7 dpi (**C**) of mock- and MAYV-infected lean and obese mice. See also Figures S4 and S5 for the cell clusters identified in each group. (**D-G**) The proportion of each cell type was identified by scRNA-seq of blood at 2dpi and 7dpi from lean (**D-E**) and obese (**F-G)** mock and MAYV-infected groups.

### Single-cell analysis of early MAYV infection reveals aberrant activation of inflammatory monocytes and reduction of B and T cell populations in lean and obese mice

To interrogate peripheral blood immune cell dynamics early during infection at peak viremia, we analyzed differences in the proportion and gene expression profiles of blood immune populations at 2 dpi. MAYV infection caused a significant drop in B cells in both lean (64.08% mock and 27.94% infected) and obese (69.43% mock and 36.57% infected) mice (Figure 3A). We also observed a modest decrease in T cells in lean (15.74% mock and 11.93% infected) and obese (9.34% mock and 9.60% infected) groups. In contrast, monocytes increased ∼4.8-fold in lean (7.96% mock and 38.24% infected) and ∼4-fold in obese (8.6% mock and 34.43% infected) mice in response to MAYV infection. Interestingly, a small population of basophils (1.29%) and stromal cells (2.11%) were detected in MAYV-infected obese mice, which were not detected in the MAYV-infected lean group (Figure 3A). It was previously shown that a subset of adipose-associated stromal cells express CD45, possibly explaining their presence in the blood of obese mice^33^.

**Figure 3.**
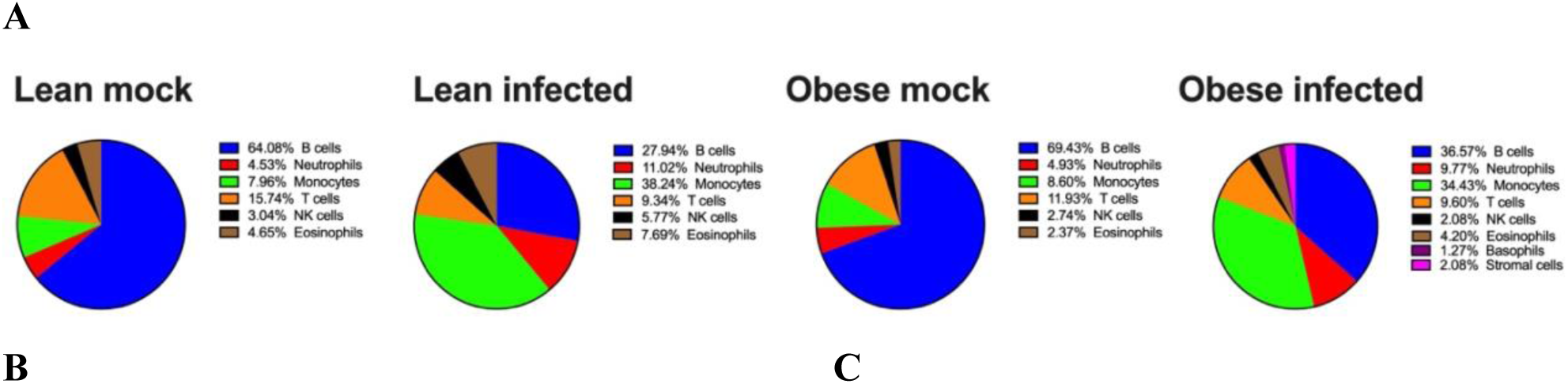
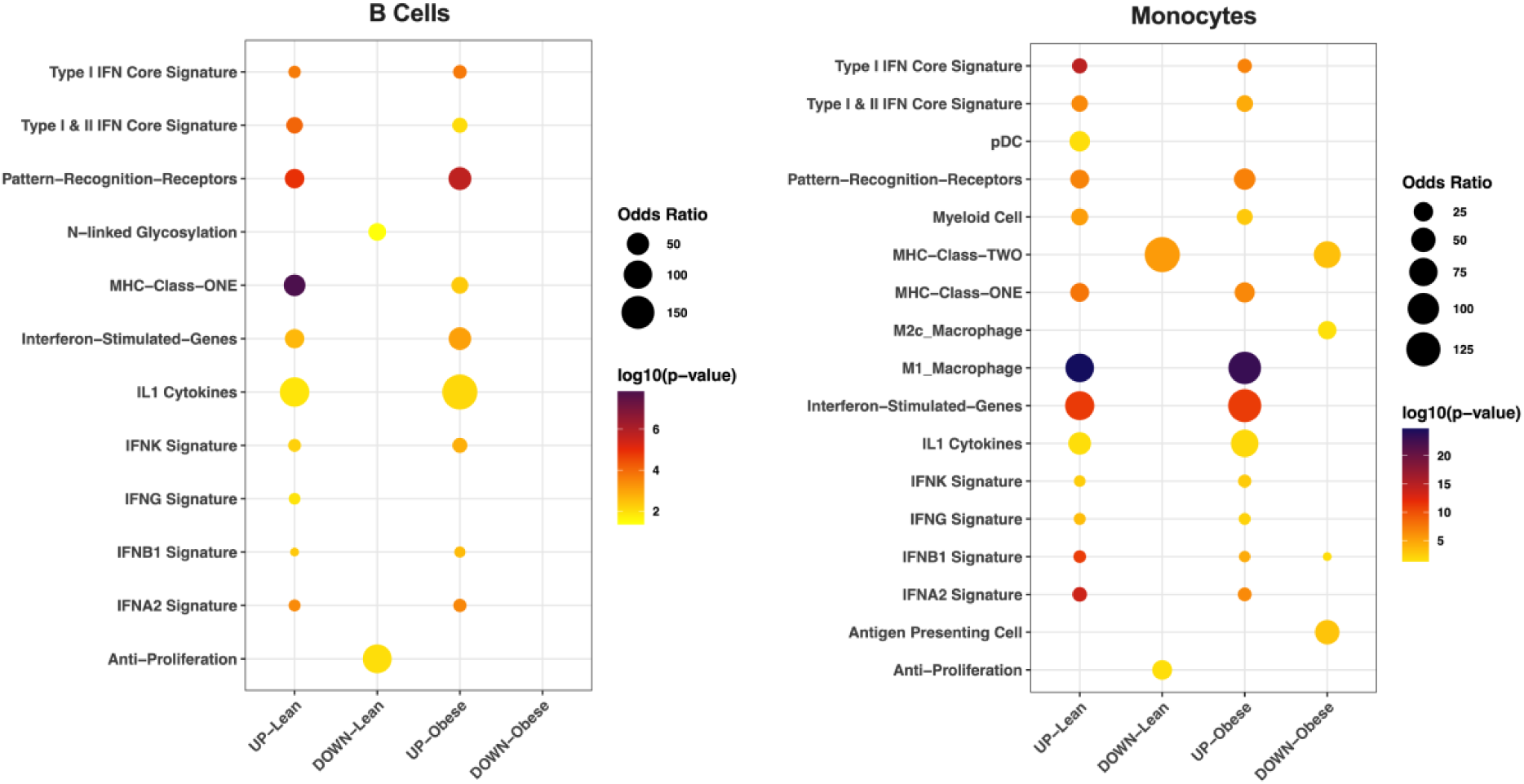
scRNA-seq during peak viremia reveals aberrant activation of inflammatory monocytes and reduction of B and T cell populations in lean and obese mice in response to MAYV infection. (**A**) Pie charts show the proportion of immune cells identified in lean and obese mock- and MAYV-infected animals at 2 dpi with scRNA-seq. (**B-C**) Dot plots depict the differential expression of genes in B cells (**B**) and monocytes (**C**) in lean and obese animals. Color change shows the p value difference in log10 compared to mock group and dot size present the odds ratio difference associated with virus infection. UP: increased expression in infected group, DOWN: decreased expression in infected group.

Next, we identified differentially expressed genes (DEGs) between mock- and MAYV-infected blood immune cell populations from lean and obese mice at 2 dpi (Figure 3B-C). For each cell population, up- and down-regulated genes in infected mice were used to identify significant overlaps with curated immune cell and pathway gene signatures. B cells and monocytes from infected mice were enriched for signatures indicative of response to type I and type II IFN, consistent with an acute response to virus exposure (Figure 3B). In addition, both B cells and monocyte populations were enriched for IL1 cytokine, MHC Class I, and pattern recognition receptor gene signatures. B cells from lean mice had a more robust enrichment of MHC Class I genes; furthermore, these cells had unique downregulation of genes in N-linked glycosylation and anti-proliferation pathways. N-linked glycosylation has been linked to B cell maturity^34^. Monocytes from infected mice also exhibited increased expression of M1 macrophage genes and decreased expression of genes involved in antigen presentation as compared to mock infection (Figure 3C). Overall, gene set enrichment was similar between lean and obese mice at 2 dpi. Thus, the initial response to arthritogenic alphavirus infection is characterized by an intense IFN response, B cell lymphopenia, and an increase in M1-like monocytes in both lean and obese mice.

### Early abnormalities in blood immune cell populations are largely resolved later during MAYV infection

We next evaluated the peripheral immune cell transcriptome at 7 dpi, when significant differences in weight loss were observed between MAYV-infected lean and obese mice (Figure 1A). The decrease in B cells in virus-infected mice observed at 2 dpi was largely restored at 7 dpi (Figure 4A). The increased proportion of monocytes at 2 dpi was no longer present at the 7 dpi. The proportions of other immune cell populations at 7 dpi in the blood were comparable between lean and obese mice (Figure 4A).

**Figure 4.**
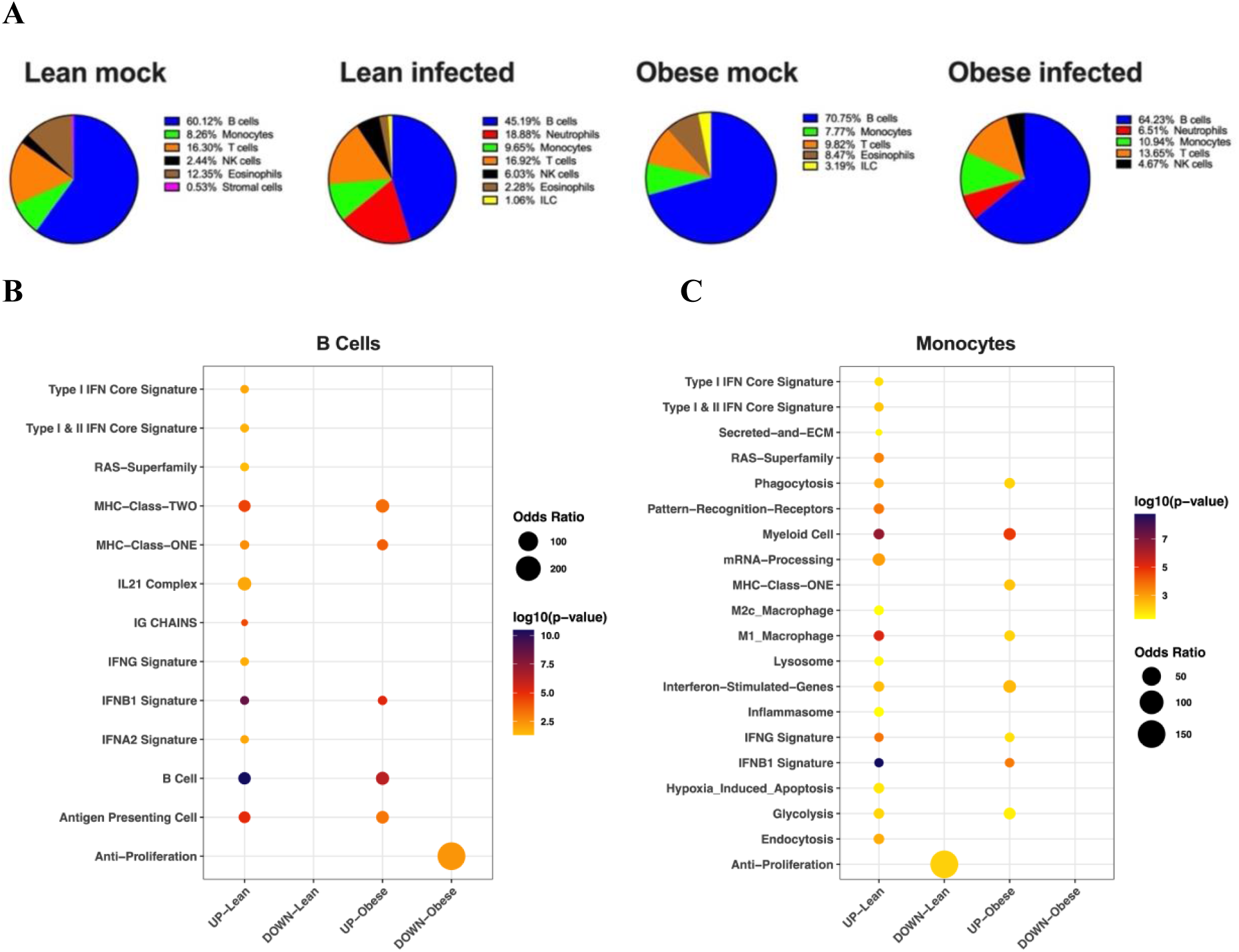
Resolution of blood abnormalities in immune cell populations late in infection. (**A**) Pie charts show the proportion of immune cells identified in the blood of lean and obese mock- and MAYV-infected mice at 7 dpi with scRNA-seq. (**B-C**) Dot plots depict the differential expression of genes in B cells (**B**) and monocytes (**C**) from lean and obese mice. Color change shows the p-value difference in log10 compared to mock group and dot size represents the odds ratio difference due to virus infection. UP: increased expression in virus infected group, DOWN: decreased expression in virus infected group.

Interestingly, changes in gene expression between lean and obese mice were more apparent at this later time point than at 2 dpi in both B cells and monocytes. B cell from infected lean mice showed enriched expression of type I and II IFN response genes and IL-21 signaling that was not observed in B cells from obese mice (Figure 4B). Similarly, we observed upregulation of IFN stimulated genes along with enriched signatures of phagocytosis, glycolysis, and signatures for both M1-and M2c-like macrophages in infected lean monocytes (Figure 4C). Enrichment for M2c-related genes at this later time point may indicate a shift towards a more anti-inflammatory state, consistent with reduced weight loss in lean mice (Figure 1A). In monocytes from infected obese mice, the enriched gene signatures were similar to those observed in lean animals; however, the magnitude of enrichment was reduced (Figure 4C). Thus, the immune response to infection at 7 dpi appears to be reduced in obese mice.

### Single-cell profiling of footpad immune cells during MAYV infection at the time of peak footpad swelling

To explore individual immune cell transcriptomes during peak footpad swelling, we collected footpads from lean and obese mock- and MAYV-infected mice at 7 dpi, isolated leukocytes and performed scRNA-seq. The UMAP of annotated cell type clusters showed an appreciable difference in macrophage and B cell clusters in response to virus infection (Figure 5A). Macrophages comprised 54.94% and 59.36% of total immune cells in lean and obese MAYV-infected animal footpads, respectively (Figure 5B). In contrast, macrophages comprised only 27.38% and 25.45% of immune cells in mock-infected lean and obese mice, respectively. Neutrophils and B cells were reduced in response to infection, whereas monocytes, dendritic cells, and T/NKT cells were increased compared to mock-infected groups in both lean and obese MAYV-infected animals (Figure 5B).

**Figure 5.**
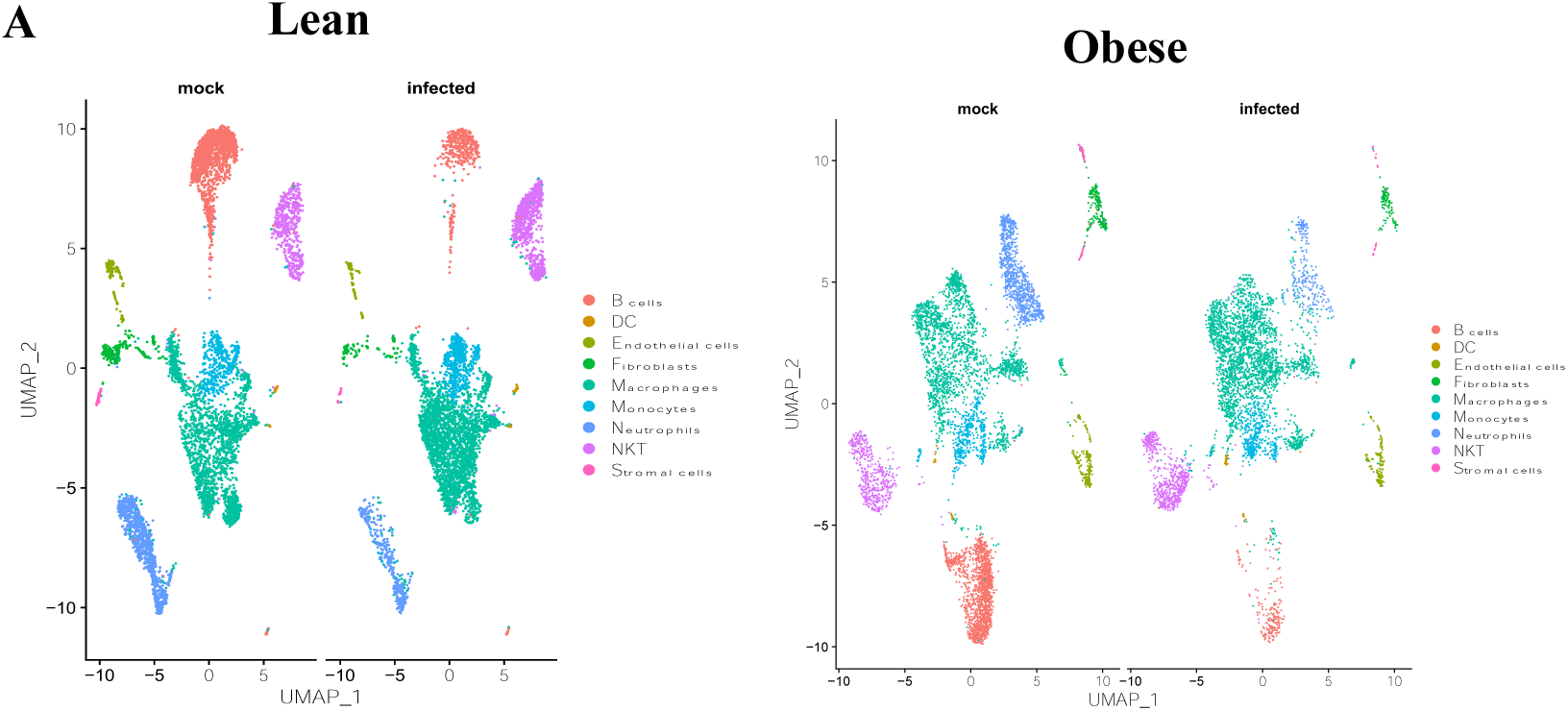
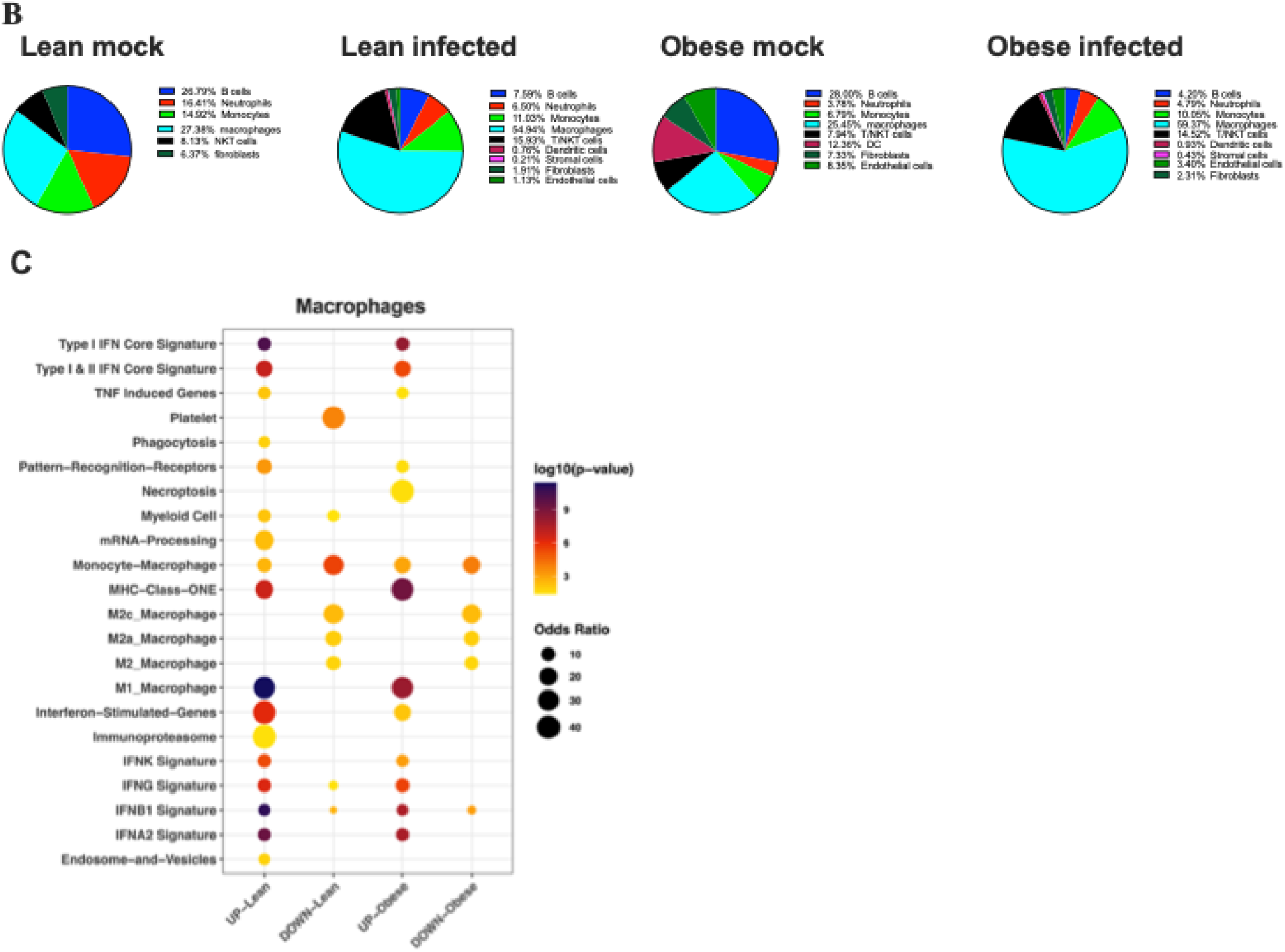
Assessment of footpad immune cells during MAYV infection at the time of peak footpad swelling. (**A**) UMAP presents the cell clusters detected in footpads at 7 dpi in MAYV-infected lean and obese animals. (**B**) Pie charts show the proportion of immune cell clusters identified in lean and obese mock- and MAYV-infected animal footpads. (**C**) Dot plots depict the differential expression of genes in lean and obese macrophages. Color change shows the p value difference in log10 compared to mock group and dot size represents the odds ratio difference due to virus infection. UP: increased expression in virus infected group, DOWN: decreased expression in virus infected group.

### Macrophages from obese mice have a unique phenotype

Macrophages are critical in arthritogenic alphavirus disease development^35^. We observed a significant increase in macrophages in footpads from lean and obese mice following MAYV infection (Figure 5B). Thus, we compared macrophage transcriptional signatures in mock- and MAYV-infected mice. MAYV infection induced similar transcriptional changes in macrophages for both lean and obese mice, including enrichment of type I and II IFN response genes, M1-like macrophage, and MHC Class I signatures and downregulation of M2-like macrophage signatures (Figure 5C). However, the IFN-stimulated gene signature, pattern-recognition receptor, IFNK, and IFNB1 gene signatures were more highly enriched in lean than obese mice (Figure 5C). Several pathways were enriched in macrophages from lean but not obese mice including mRNA processing, immunoproteasome, phagocytosis, and endosome and vesicles (Figure 5C). Necroptosis-related pathways were only upregulated in obese macrophages.

### Unique transcriptional signatures define the dominant macrophage populations in the footpads of lean and obese mice during MAYV infection

We next evaluated specific subsets of macrophages in the footpads of lean and obese mice. Each subset was annotated based on transcriptional overlaps with previously defined macrophage populations from the ImmGen database^36^. We identified 5 macrophage subsets in lean mock- and MAYV-infected footpads: MF.103-11B+SALM3, MF.103-11B+, MF.103CLOSER, MF.F480HI.GATA6KO, and MFIO5.II+480LO (Figure 6A, Table 1). Whereas the frequency of 4/5 of the macrophage subsets was reduced in MAYV-vs. mock-infected animals, the MFIO5.II+480LO subset increased from 39.15% in mock to 58.25% in MAYV-infected mice. Eight distinct macrophage subclusters were detected in mock- and MAYV-infected obese mice: MF.103-11B+, MF.103-11B+24-, MF.103CLOSER, MF.F480HI.CTRL, MF.F480HI.GATA6KO, MF.RP, MFIO5.II-480HI, and MFIO5.II+480INT (Figure 6A, Table 2). Similar to lean mice, 6/8 macrophage populations were reduced in MAYV-vs. mock-infected obese mice; however, MFIO5.II+480INT (mock 47.37%; infected 70.15%) increased in response to infection (Table 2). The dominant macrophage subset was MFIO5.II+480LO and MFIO5.II+480INT in lean and obese mice, respectively (Table 1 and 2). This shift from F480^lo^ macrophages in lean mice to F480^int^ in obese mice suggests a less activated macrophage population in obese animals (Figure 6A, Table 1 and 2).

**Figure 6.**
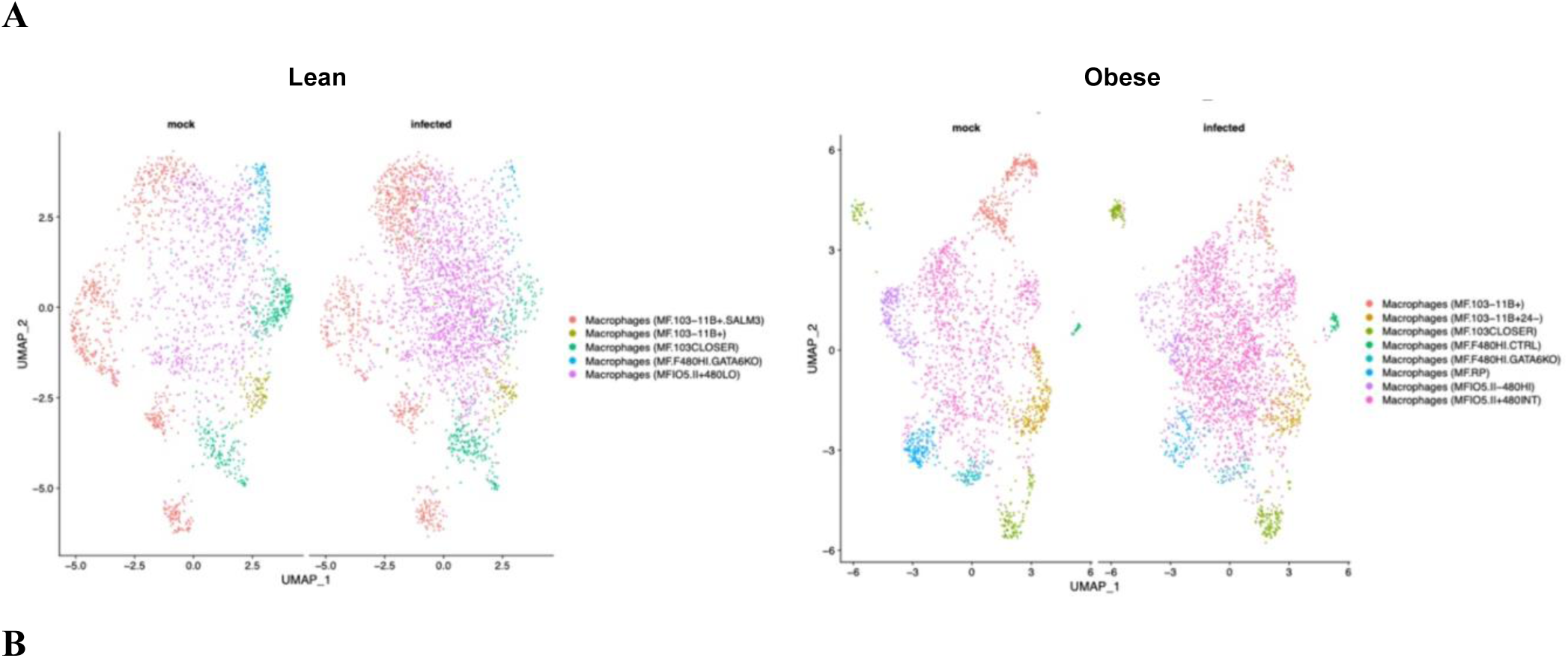
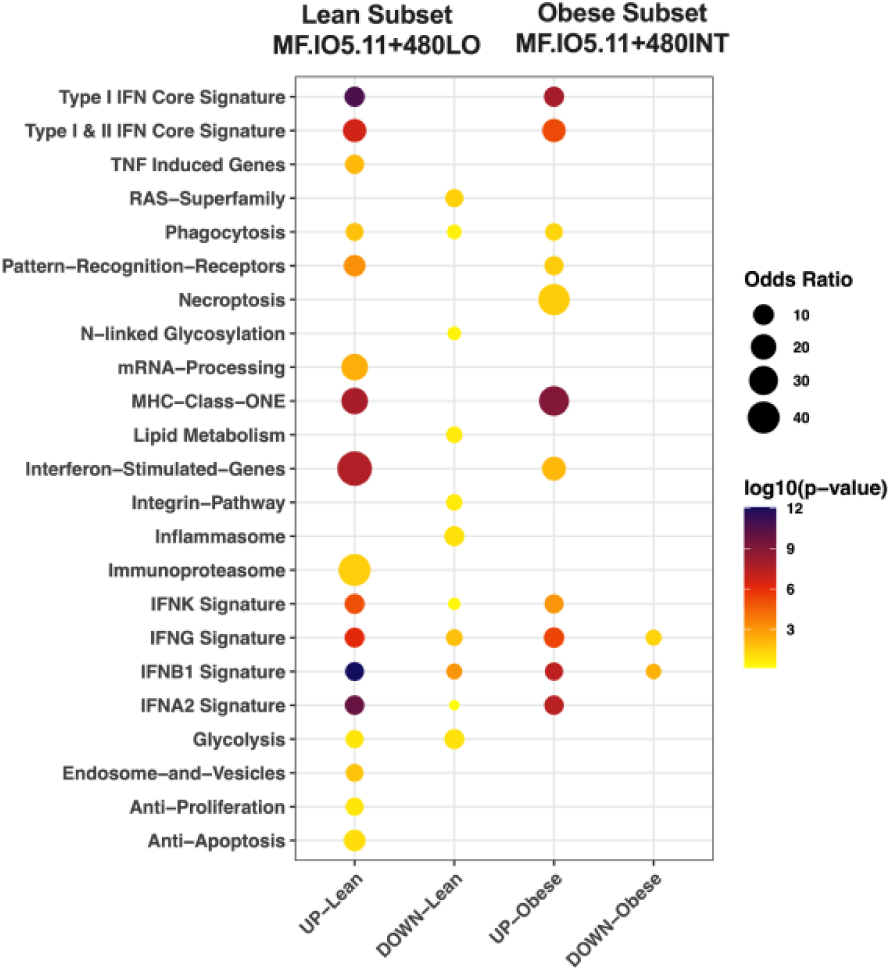
Dynamic shifts in macrophage populations in lean and obese mice during MAYV infection. Macrophage populations identified in mock and MAYV-infected lean and obese animal footpads were further analyzed to explore the different subclusters of macrophages raised in response to MAYV infection and their transcriptional profiles. (**A**) Macrophage subclusters detected in lean and obese mock- and MAYV-infected animal footpads at 7 dpi. (**B**) Lean macrophage subset CSF1R^+^SiglecF^-^F4/80^+LOW^MHCII^+high^ and obese macrophage subset CSF1R^+^SiglecF^-^F4/80^+INT^MHCII^+high^ transcriptional profiles detected in MAYV-infected lean and obese animals compared to mock groups. Color change shows the p-value difference in log10 compared to the mock group, and dot size represents the odds ratio difference due to virus infection. UP: increased expression in virus infected group, DOWN: decreased expression in virus infected group.

**Table 1:**
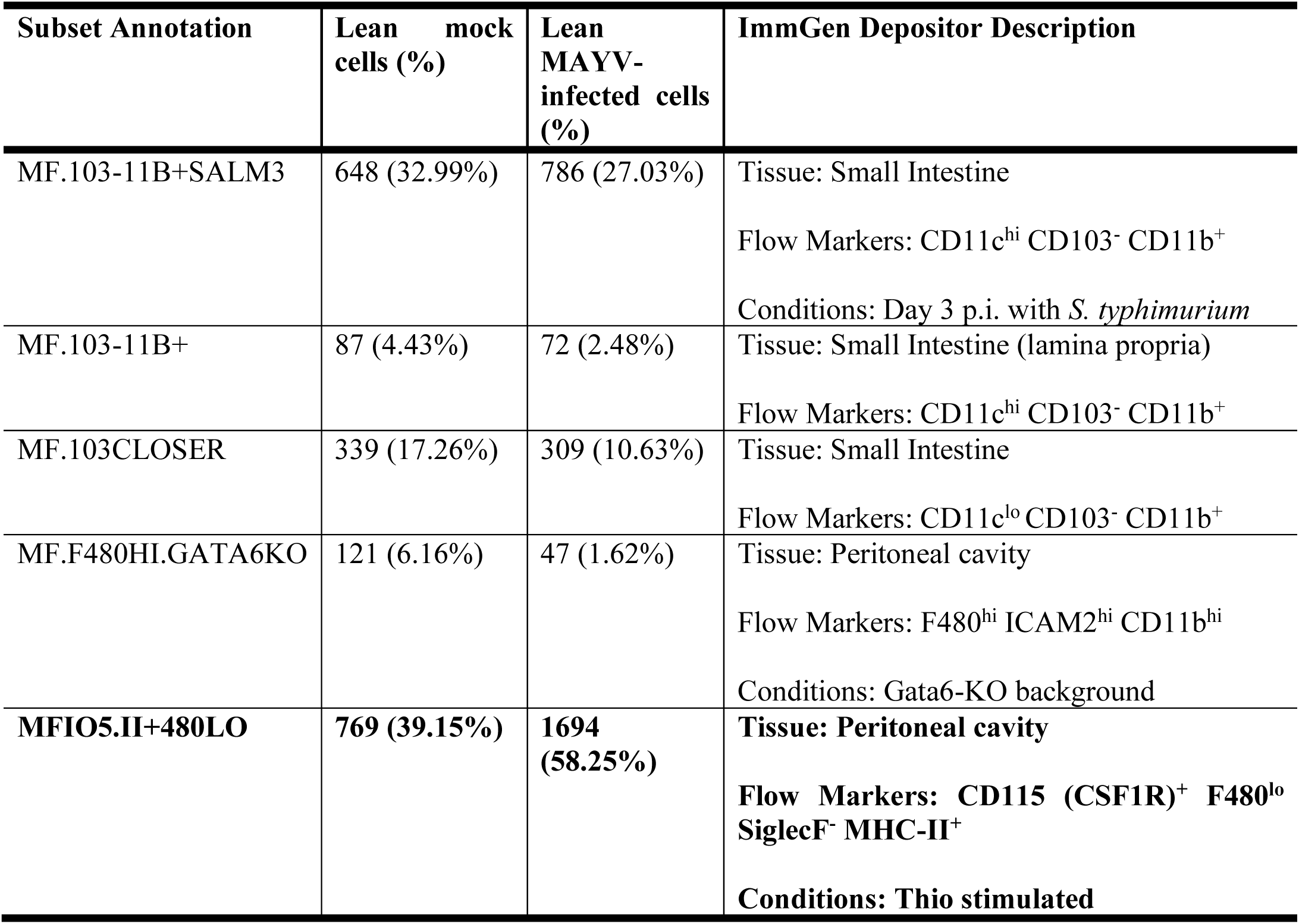
Macrophage subsets identified in lean mock- and MAYV-infected animal footpads.

| Subset Annotation | Lean mock cells (%) | Lean MAYV-infected cells (%) | ImmGen Depositor Description |
| --- | --- | --- | --- |
| MF.103-11B+SALM3 | 648 (32.99%) | 786 (27.03%) | Tissue: Small Intestine<br>Flow Markers: CD11c<sup>hi</sup> CD103<sup>-</sup> CD11b<sup>+</sup><br>Conditions: Day 3 p.i. with *S. typhimurium* |
| MF.103-11B+ | 87 (4.43%) | 72 (2.48%) | Tissue: Small Intestine (lamina propria)<br>Flow Markers: CD11c<sup>hi</sup> CD103<sup>-</sup> CD11b<sup>+</sup> |
| MF.103CLOSER | 339 (17.26%) | 309 (10.63%) | Tissue: Small Intestine<br>Flow Markers: CD11c<sup>lo</sup> CD103<sup>-</sup> CD11b<sup>+</sup> |
| MF.F480HI.GATA6KO | 121 (6.16%) | 47 (1.62%) | Tissue: Peritoneal cavity<br>Flow Markers: F480<sup>hi</sup> ICAM2<sup>hi</sup> CD11b<sup>hi</sup><br>Conditions: Gata6-KO background |
| MFIO5.II+480LO | 769 (39.15%) | 1694 (58.25%) | **Tissue: Peritoneal cavity**<br>**Flow Markers: CD115 (CSF1R)<sup>+</sup> F480<sup>lo</sup> SiglecF<sup>-</sup> MHC-II<sup>+</sup>**<br>**Conditions: Thio stimulated** |

**Table 2:**
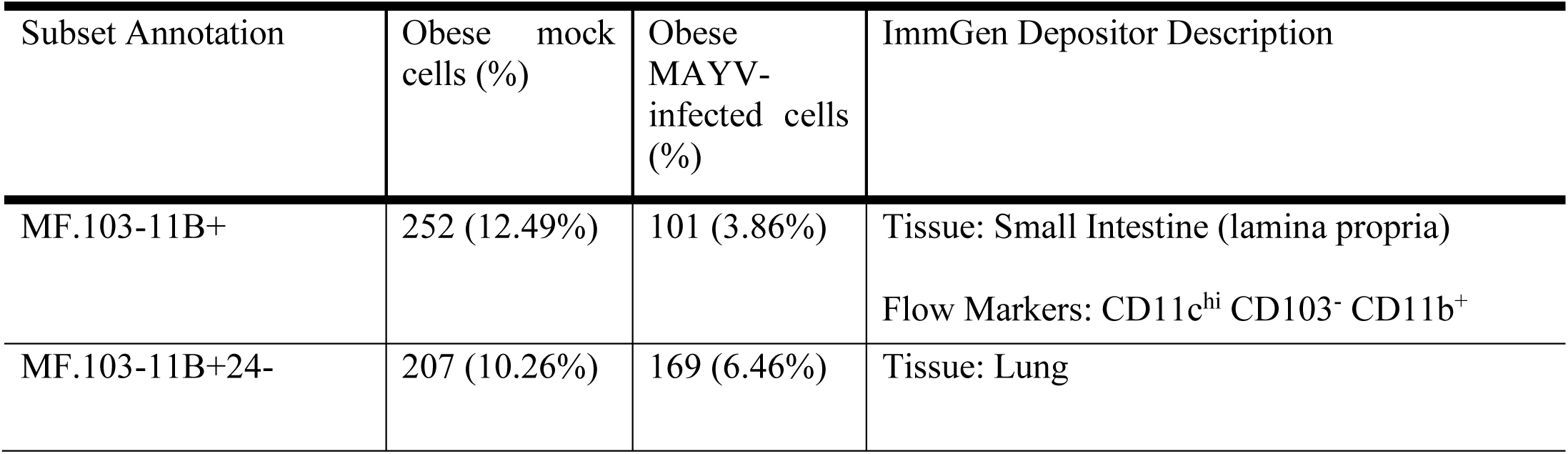
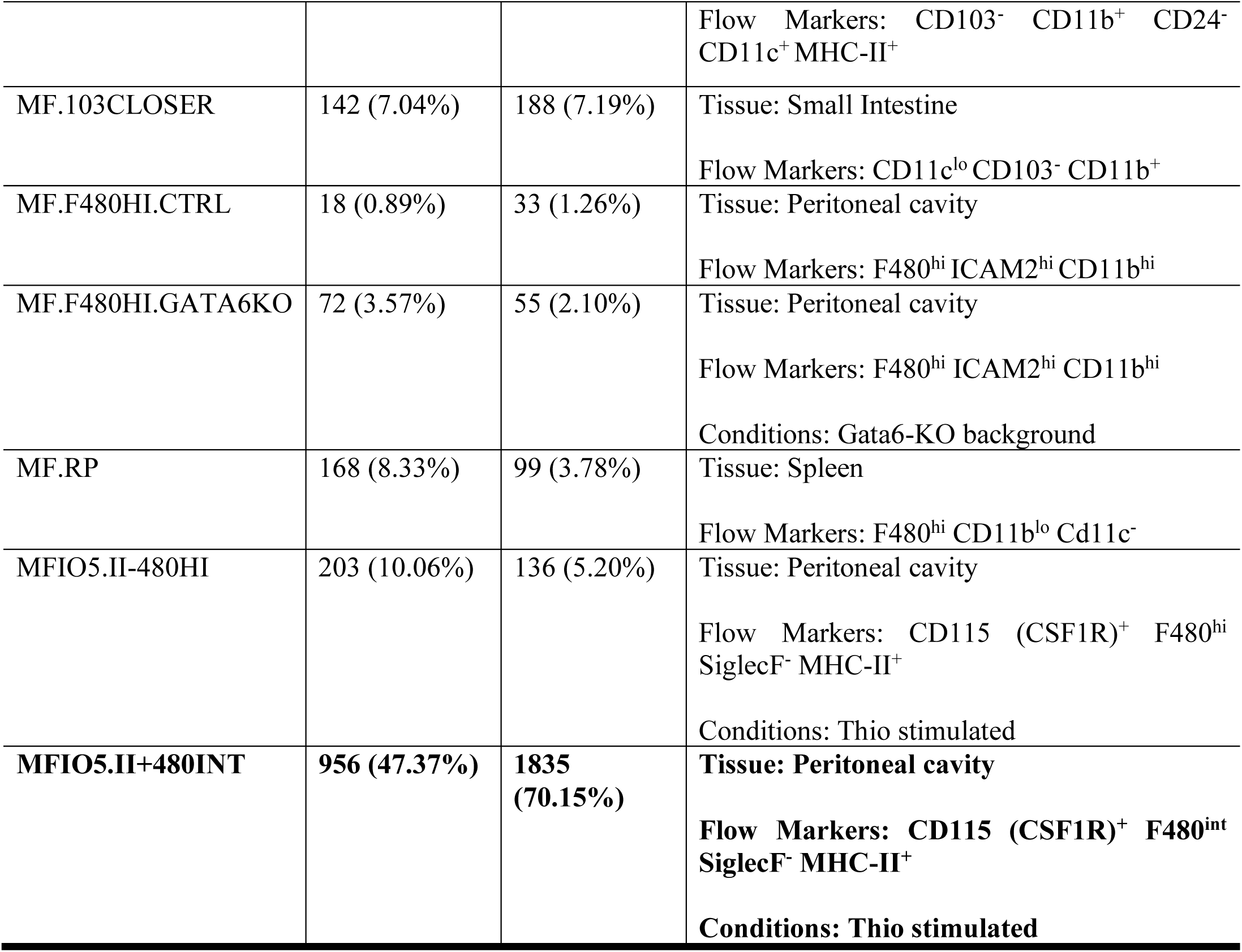
Macrophage subsets identified in obese mock- and MAYV-infected animal footpads.

To test this hypothesis, we analyzed the transcriptomes of lean MFIO5.II+480LO (CSF1R^+^SiglecF^-^F4/80^+LOW^MHCII^+high^) and obese MFIO5.II+480INT (CSF1R^+^SiglecF^-^ F4/80^+INT^MHCII^+high^) macrophage populations. We observed that the lean mice macrophage subset was enriched for TNF-induced genes, mRNA processing, immunoproteasome, glycolysis, endosome and vesicle, anti-proliferation, and anti-apoptosis gene signatures, and expressed a robust IFN response signature, which was not observed in the obese macrophage subset (Figure 6B). Moreover, N-linked glycosylation, RAS superfamily, lipid metabolism, and integrin pathways were uniquely downregulated in the lean macrophage subset. In contrast, the necroptosis signature was uniquely enriched in the obese macrophage population.

## Discussion

In 2022, more than 800 million people were considered obese (BMI > 30) worldwide^10^, and these numbers are continuing to rise. This is concerning given the health risks associated with the development of obesity and the higher risk of severe disease following infection with different viruses, including arthritogenic alphaviruses^8,37^. Arthritogenic alphavirus infection induces footpad swelling, myositis, and bone loss in mouse models, similar to human disease^11,21–23^. Here, we confirmed that obesity leads to worse disease outcomes following arthritogenic alphavirus infection, including increased weight loss and worse tissue inflammation in obese compared to lean mice. Toward understanding the mechanisms underlying these differences, we performed a comprehensive profiling of immune cell phenotypes at various stages of arthritogenic alphavirus disease development in lean and obese mice using scRNA-seq. We observed large shifts in immune cell populations in the blood and footpads of lean and obese mice infected with MAYV. Infection was associated with significant enrichment of genes in known antiviral signaling pathways, along with shifts in antigen presentation, cytokine signaling, and cell death-related genes. In the footpad, we detected lower expression of antiviral gene signatures across most cell types in obese compared to lean mice. Furthermore, we identified a unique CSF1R^+^SiglecF^-^ F4/80loMHCII^+high^ macrophage subset in lean mice with enhanced expression of IFN response genes that was not found in obese mice and may account for the diminished pathogenic consequences of MAYV infection in lean mice.

We inoculated lean and obese mice with MAYV and found that obese animals developed more severe disease in line with our previous reports showing that infection with CHIKV, MAYV, or RRV led to worse disease outcomes in obese mice^11^. To evaluate immune cell dynamics during MAYV infection, we carried out scRNAseq at two time points: at 2 dpi, peak viremia^38^, and 7 dpi, peak footpad swelling. At 2 dpi, we observed a significant drop in B cells and an increase in monocytes in lean and obese mice. A similar trend was observed in previous reports of CHIKV-infected human blood cells^25^. It has been reported that type I IFN-dependent signaling leads to early suppression of humoral responses when virus-infected B cells are eliminated by inflammatory monocytes^39^ and CD8 T cells^40^. Type I IFN-related gene pathways were significantly upregulated in monocytes, and it has been reported that blocking type I IFN can result in restoration of the B cell response to the virus during acute infection^41^. A direct role of B cells in mediating the resolution of arthritogenic alphavirus disease has also been reported^42,43^. CHIKV infection in B cell-deficient mice led to persistent infection in the joint, and disease resolution was dependent on antibody-mediated neutralizing activity, cellular cytotoxicity, and complement activation^43,44^. CHIKV infection in human B cells has been shown in human patient blood cells^25^ and *in vitro* experiments^45^. Interestingly, our B cell transcriptomic data showed that MHC class I signaling genes were highly upregulated in B cells from both lean and obese hosts at 2 dpi, possibly indicative of viral presentation by infected B cells to cytotoxic T cells. The potential relationship between a decrease in B cells and the development of more severe disease should be explored in future studies.

Blood monocytes increased significantly in response to infection in both lean and obese mice at 2 dpi. Recently, it was reported that Ly6C^+^ monocytes facilitate alphavirus infection at the initial infection site, which promotes more rapid spread into circulation^46^. Furthermore, monocyte recruitment to the draining lymph nodes during CHIKV infection impairs virus-specific B cell responses by virtue of their ability to produce nitric oxide^47^. The substantial reduction in B cells, along with a concomitant rise in monocytes following infection should be explored in future studies.

Arthritogenic alphavirus infection in mice produces significant weight loss and footpad swelling at 7 dpi^11^. We collected blood and footpad immune cells at 7 dpi to determine the effect of infection on immune cell populations during later stages of infection. Most of the cell populations recovered in infected lean and obese mouse blood returned to baseline levels. While we observed only minor differences in immune cell percentages between infected lean and obese animals, changes in gene expression were more apparent at 7 dpi compared to 2 dpi. B cells from lean mice showed enriched expression of type I and II IFN signature genes, IFNG, IFNB1, IFNA2 response genes, and IL-21 signaling. B cells from obese mice showed enrichment of genes associated with IFNB1 exposure but none of the other antiviral signaling gene sets and the IL-21 pathway remained unchanged. Higher expression of MHC II-related genes in lean B cells is likely indicative of their role as antigen-presenting cells^48,49^. Enrichment of IL-21 signaling may explain the higher proliferation and activation of B cells at 7 dpi in lean compared to obese mice, which is predominately mediated through IL-21^50^. Furthermore, we observed upregulation of IFN-associated signaling pathways along with enriched signatures of mRNA processing, endocytosis, and M2c-like gene sets in monocytes from lean compared to obese mice. Enrichment for M2c related genes at this later time point may indicate a shift towards a more anti-inflammatory state in lean mice^51,52^. Worsened systemic disease outcomes in obese mice could be explained by the dampened expression of type I and II signature genes, IFNG, IFNB1, and IFNA2 response signatures in blood B cells and monocytes.

We observed significantly greater footpad swelling and tissue damage in obese compared to lean mice. During alphavirus infection, chemokines attract monocytes and other immune cells to the site of infection^53–57^. The monocytes differentiate into macrophages in infected tissues and secrete proinflammatory cytokines such as TNF, IL-6, IL-1β, and type I IFNs^45,58–60^. Arthritogenic alphaviruses replicate in monocytes and tissue resident macrophages and may contribute to chronic disease^35,45,61,62^. scRNA-seq revealed a considerable increase in macrophages and T/NKT cells at 7 dpi in the footpads of lean and obese mice. In contrast, B cells dropped significantly in response to infection. Flow cytometric validation also revealed that CD4^+^ T cells were significantly higher in obese compared to lean mice following infection. Previous data shows that CD4^+^ T cells contribute to the development of pathological damage during arthritogenic alphavirus infection^63,64^, suggesting these cells may contribute to worse tissue damage observed in obese mice.

Interestingly, M2c, M2a, and M2 macrophage gene expression was significantly reduced in MAYV-infected lean and obese footpad macrophages, which may highlight the loss of anti-inflammatory mediators^65^ during peak footpad swelling. However, IFN stimulated gene signatures were more highly enriched in lean macrophages. IFN response genes are involved in viral sensing and mediating antiviral, immunomodulatory, and antiproliferative effects^66,67^; thus, the reduced expression in obese host suggests an impaired immune response to MAYV infection. Consistent with this, we previously observed higher virus replication in the footpad of obese mice at peak footpad swelling compared to lean mice ^11^. Several IFN response genes with important roles in viral response were uniquely enriched in macrophages from lean infected mice. Similarly, mRNA processing genes activated in lean macrophages are involved in higher synthesis, modification, mRNA splicing, polyadenylation, and expression of ISGs^68,69^. The immunoproteasome plays a role in antigen processing and presentation^70^ and genes involved in the transcription of immunoproteasome subunits may increase antigenic peptide generation for MHC I presentation and regulation of immune responses in macrophages^71,72^. Phagocytosis and endosome and vesicle related genes encode proteins that support macrophage functions to engulf cellular debris and help in clearance of virus infected cells and control of virus replication^73–81^. The enrichment of these pathways in lean hosts suggests a superior response to virus infection, which may result in better control of virus replication and subsequent disease. In contrast, the unique upregulation of necroptosis-related genes in obese macrophages may contribute to the increase tissue damage observed. Its selective upregulation in obese macrophages suggests a potential link between obesity-related metabolic dysregulation and inflammatory cell death processes, which requires further investigation.

We also carried out macrophage subset analysis and found two main subclusters: MFIO5.II+480^LO^ and MFIO5.II+480^INT^ in lean and obese mice, respectively. F4/80 is a cell surface marker for murine macrophages, which is expressed on resident tissue macrophages and is associated with maturation status^74,75^. Previous studies highlight that F4/80 expression is decreased in activated macrophages as they engulf viral particles and virus infected cells, and process viral proteins to present through MHC-II receptors^76–78^. Transcriptomic analysis of lean CSF1R^+^SiglecF^-^ F4/80^+LOW^MHCII^+high^ and obese CSF1R^+^SiglecF^-^F4/80^+INT^MHCII^+high^ macrophage populations showed that the lean macrophage subset was enriched not only in IFN response genes, but also in TNF-induced genes, mRNA processing, immunoproteasome, glycolysis, endosome and vesicle, anti-proliferation, and anti-apoptosis gene sets, that were not observed in obese macrophage subset. The upregulation of these pathways is indicative of a inferior macrophage response in obese infected mice that provides a plausible mechanism for increased tissue inflammation and footpad swelling in response to MAYV infection in obese as compared to lean hosts.

## Limitations

Our study has several limitations: (1) We used only MAYV for scRNA-seq studies, (2) performed positive selection for CD45 cells and (3) tested footpads at only a single timepoint. Thus, we could not assess the differences in response at different timepoints or the effects on most non-immune cells. Furthermore, we could not perform a comprehensive comparison between arthritogenic alphaviruses. We used MAYV since it is a BSL2 virus and replicates better in mice than CHIKV; however, future studies should provide a more comprehensive comparison between viruses at several timepoints following infection, including during the chronic stage.

## Summary

Our analysis provides a detailed profiling, at the single cell level, of the lean and obese host immune response to arthritogenic alphavirus infection. These studies enhance our understanding of the impact of obesity on immune cell dynamics during viral infection. This work provides multiple targets for therapeutic development, which could prove vital for combating arthritogenic alphavirus disease. The described macrophage subsets and related pathways can be studied further to understand the immunopathogenesis of arthritogenic alphavirus disease and to select targets for therapeutic development. Overall, this study highlights that obesity alters immune cell functional dynamics response to arthritogenic alphavirus infection, which is associated with worse disease outcomes.

## Acknowledgments

We are grateful to Andria Doty, Colleen Palmateer, and Jason Monroe for assistance in scRNA-seq sample preparation and analysis. We also thank Melissa Makris for assisting with flow cytometry analysis. This work was funded by the NIH grant R21AI153919-01 awarded to J.W-L. The graphical abstract and figures were created using BioRender (https://biorender.com).

## Author contributions

M.H. and J.W.-L. have prepared the experiment plan and wrote the manuscript. M.H. and M.S.H performed experiments. A.D. and P.L. analyzed scRNA-seq data and assisted with manuscript preparation. S.C-O. analyzed histopathology.

## Supplementary figure legends

**Supplementary figure 1. Cytokine expression in before and after MAYV infection in lean and obese mice.** Mice were bled prior to or 2 days post-MAYV infection and cytokines were measured by multiplex Luminex assay. Comparisons were made using unpaired t tests; *p<0.05, **p<0.01, ***p<0.001, ****p<0.0001.

**Supplementary figure 2. Antiviral gene expression in lean and obese MAYV infected mice footpads.**

C57BL/6J lean and obese mice were infected with 10^4^ PFU of MAYV strain TRVL 4675 in each hind footpad and footpads were collected at 7 days post-inoculation. RNA was extracted and subjected to RT-qPCR. Data were normalized to GAPDH and analyzed using unpaired t-tests with Welch’s correction. The error bars represent the standard deviation and bars indicate mean values.

**Supplementary figure 3. Blood immune cell clusters detected at 2 dpi from mock- and MAYV-infected lean and obese mice.**

**Supplementary figure 4. Blood immune cell clusters detected at 7 dpi from mock- and MAYV-infected lean and obese mice.**

**Supplementary table 1. List of RT-qPCR primers used to assess gene expression.**

## Lead contact

Further information and requests for resources and reagents should be directed to and will be fulfilled by the lead contact, James Weger-Lucarelli.

## Materials availability

This study did not generate new unique reagents.

## STAR Methods

### KEY RESOURCES TABLE

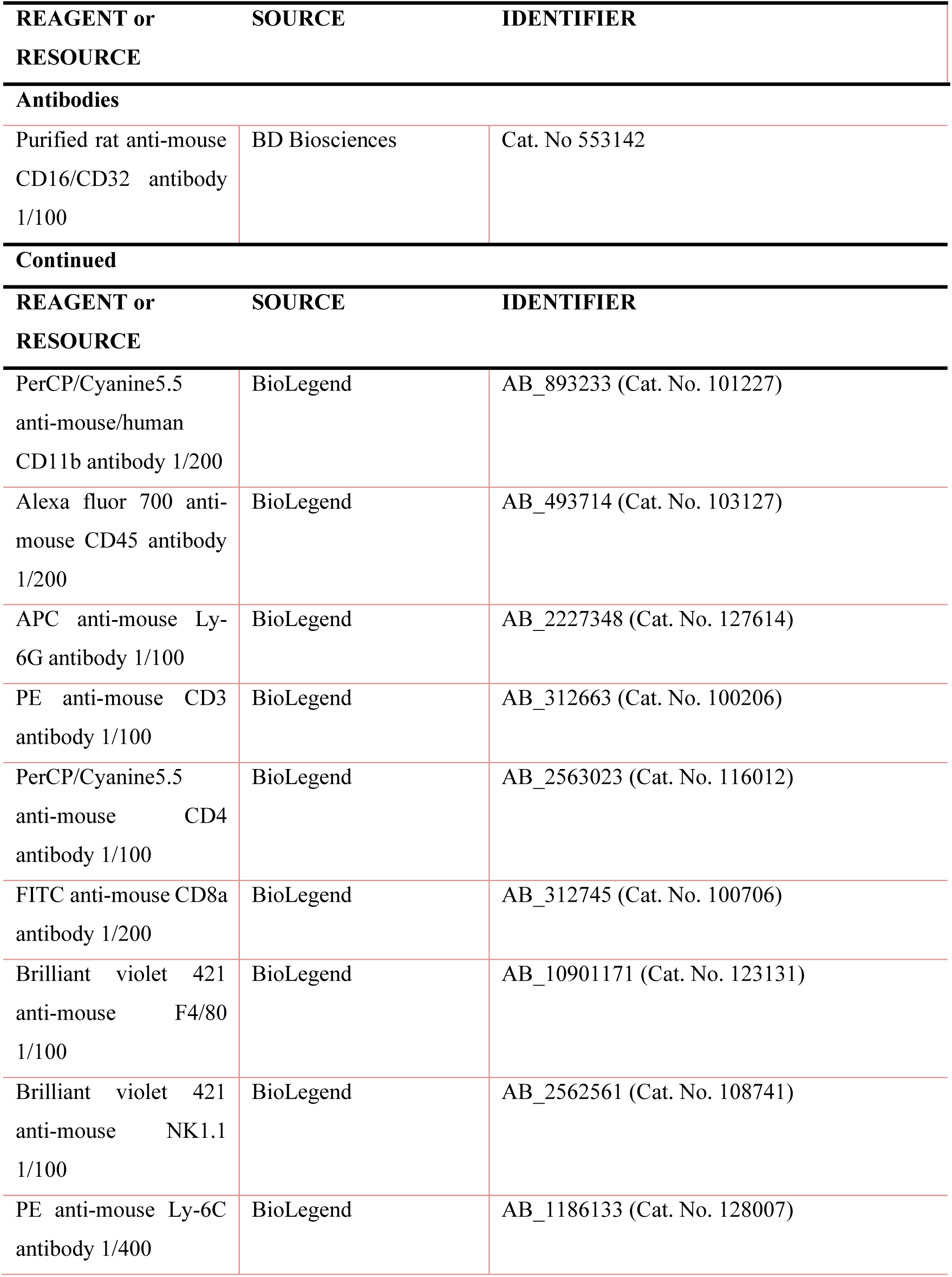
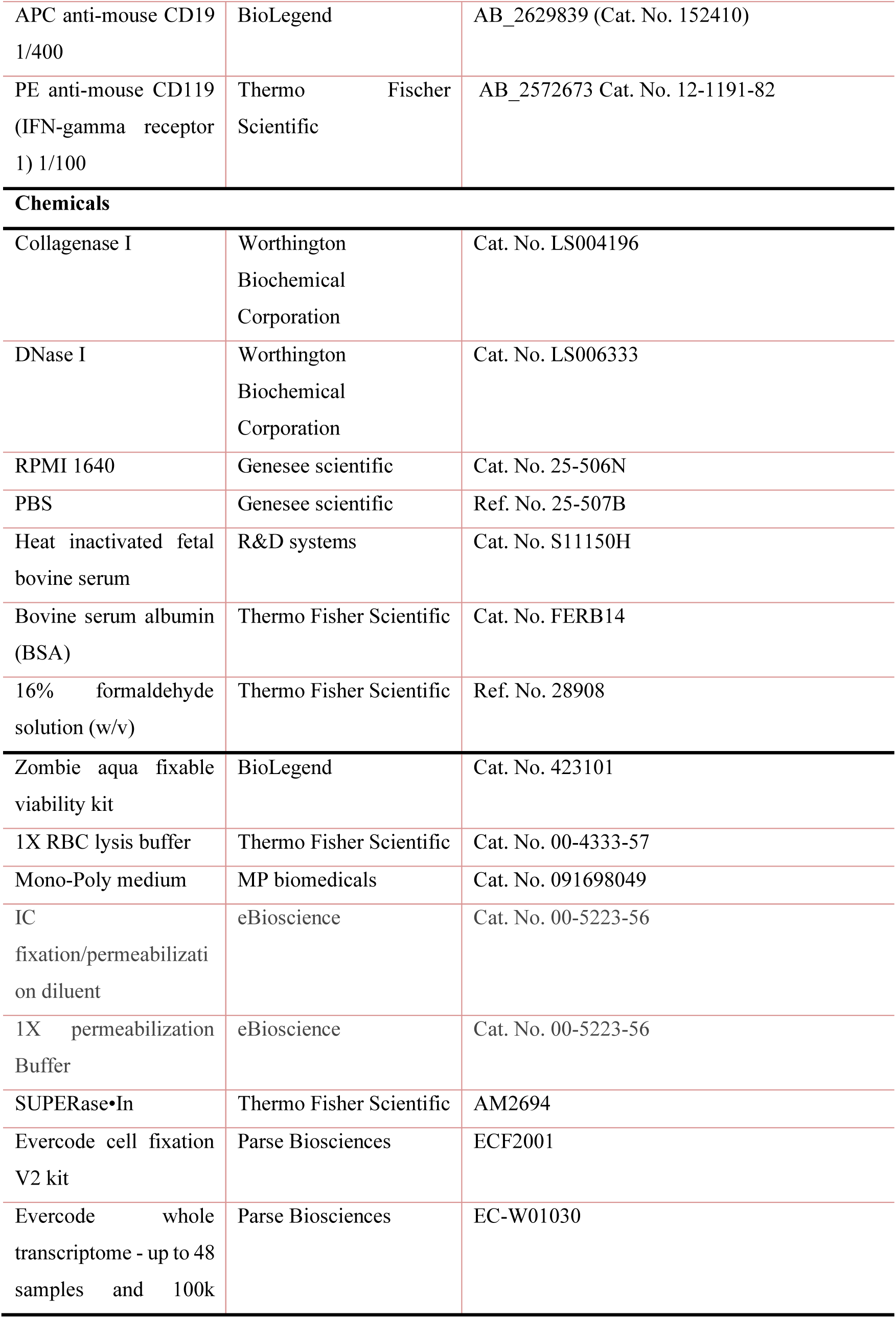
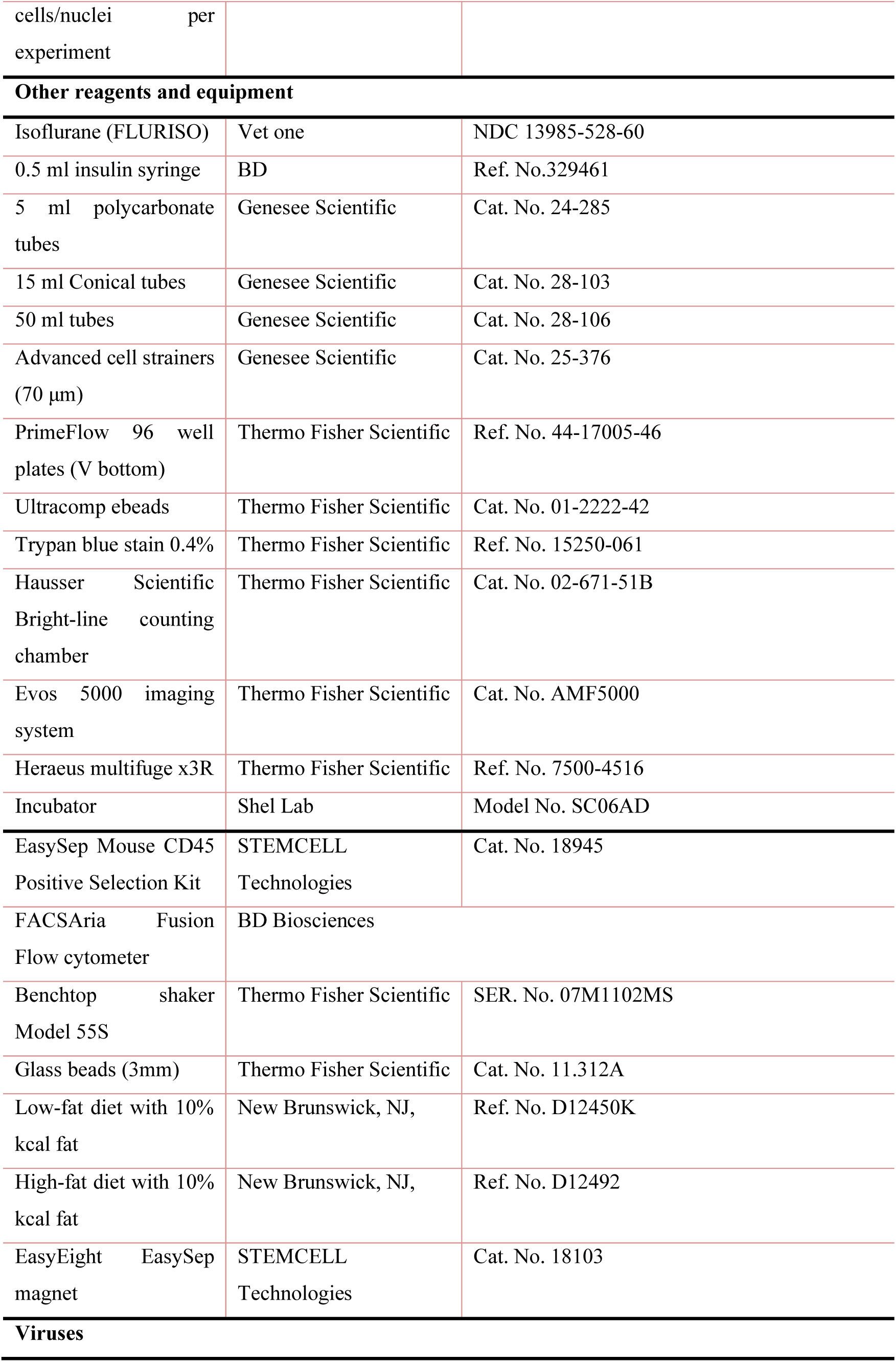
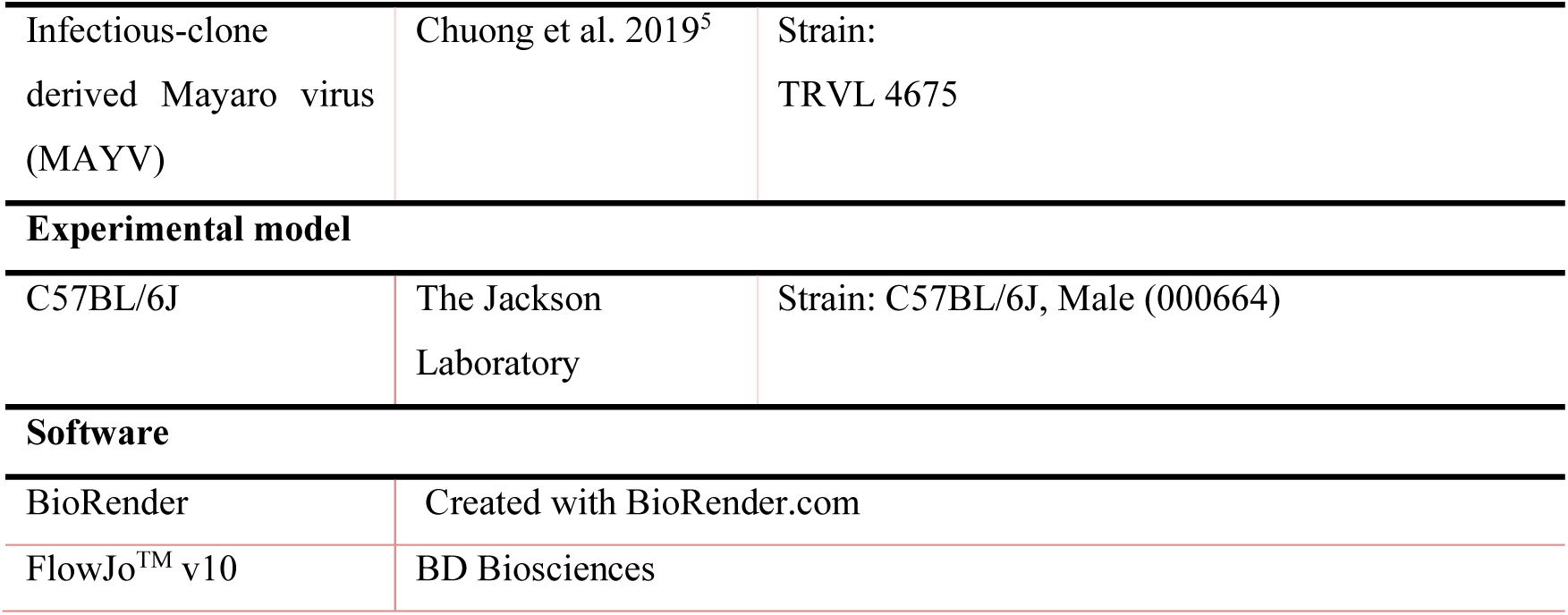

### EXPERIMENTAL MODEL AND SUBJECT DETAILS

#### Animals and Ethics

We purchased wild-type (WT) C57BL/6J male mice from The Jackson Laboratory. Virus infections were performed in mice anesthetized with isoflurane inhalation and all efforts were made to minimize animal suffering. All experiments were performed with the approval of Virginia Tech’s Institutional Animal Care & Use Committee (IACUC) under protocol number 24-060.

#### Cell lines

Vero cells were obtained from American Type Culture Collection and cultured at 37 °C in Dulbecco’s Modified Eagle Medium (DMEM) supplemented with 5% fetal bovine serum (FBS), 1% penicillin/streptomycin, and 25mM HEPES in 5% CO_2_ incubator.

#### Viruses

MAYV strain TRVL 4675 was produced from an infectious clone that we previously described^38^. The MAYV stock was propagated on Vero cells, and supernatants were harvested at 72 h post infection, titrated by plaque assay on Vero cells and stored in aliquots at -80 °C.

#### Mouse experiments

C57BL/6N mice were housed in groups of five per cage and maintained at ambient temperature with ad libitum supply of food and water. All diets used for the study were obtained from Research Diets (New Brunswick, NJ, USA). In all studies, male mice were used and fed on a low-fat diet with 10% kcal fat and on a high-fat diet with 60% kcal fat. Throughout the manuscript, we refer to the groups as lean (low-fat diet) or obese (high-fat diet). The mice were kept on these diets for 18–20 weeks before infections, and the same diets were continued until the end of the experiment. Mice were inoculated with 10^4^ PFU of MAYV in a volume of 50 μl RPMI-1640 media through subcutaneous injections in both hind feet^11,21^. Virus dilutions were made in RPMI-1640 media supplemented with 10 mM HEPES and 1% FBS. All mouse experiments were performed under ABSL 2/3 conditions. Mice were weighed daily following infection and footpad swelling was measured daily using a digital caliper. Blood was collected from the submandibular vein with a 5 mm lancet (Goldenrod) into a purple top microtainer tube (Fisher Scientific) for immune cell isolation. Virus titers were measured by plaque assay using Vero cells.

#### Tissue processing and H&E staining

At 7 dpi, animals were euthanized, and footpads were collected and fixed in 4% formalin for 48 h. After fixation, footpads were decalcified in 10% EDTA solution at 4 °C for 2 weeks. The Virginia Tech Animal Laboratory Services (VITALS) performed paraffin embedding, sectioning, and staining with hematoxylin and eosin, and a board-certified anatomic pathologist read the slides.

#### Luminex assay

We quantified cytokines levels in the serum at pre- and post-infection from lean and obese animals using the mouse Luminex XL cytokine assay (bio-techne) according to the manufacturer’s instructions. The standard curve was generated using the optical density values of the standards, which were used to calculate the cytokine levels in each sample.

#### Isolation of blood and footpad immune cells using Mono-Poly medium

Blood was collected at 2 dpi (peak viremia) and 7 dpi (severe disease symptoms), and footpads at 7 dpi when animals developed peak footpad swelling. We isolated total immune cells using Mono-Poly (MP) medium from blood and footpads. The detailed protocol for mouse footpad digestion, leukocyte isolation, purification using CD45+ selection kit was previously described ^79^. For blood cell isolation, an equal amount of blood was pooled from five animals. Then, we layered blood onto the MP medium (1:1 MP medium:blood ratio, place tube at a slight vertical angle and slowly add blood). Centrifuge at 300 x g for 30 minutes in a swinging bucket rotor at room temperature (20-25 °C). Collect cell layers between RBC’s and plasma to isolate mononuclear and polymorphonuclear cells and add to a 15 mL conical tube containing 10 mL cold RPMI media supplemented with 10% FBS. Centrifuge at 500 x g for 10 minutes at 4 °C. Resuspend cell pellets in 1 mL RPMI media and proceed for CD45+ purification by following step by step protocol ^79^. For scRNA-seq, we pooled cells from five mice into a single sample, however, we isolated blood and footpad immune cells from each animal and processed separately for flow cytometry.

#### Flow cytometry

Single cell suspensions isolated from lean and obese mice blood and footpads were washed with phosphate buffered saline (PBS) and resuspended in 100 μL Zombie aqua dye solution (1:400 prepared in PBS) and incubated at room temperature for 15-30 minutes. 200 μL flow cytometry staining (FACS) buffer (PBS containing 2% FBS) was added and centrifuged for 5 min at 4° C. The resulting cells were resuspended in FACS buffer with 0.5 mg/mL rat anti-mouse CD16/CD32 Fc block and incubated for 15 min on ice to block Fc receptors. Combined antibody solution prepared in FACS buffer with fluorophore-conjugated antibodies presented in key resources table. 100 μL antibody cocktail was added to the single cell suspension, mixed, and incubated for 30 min on ice. Cells were washed with FACS buffer twice and 100 μL 4% formalin were added to fix cells. After 15 min incubation at room temperature, cells were washed with FACS buffer, resuspended in 100-200 μL PBS and covered with aluminum foil and proceeded for analysis. Single color controls were run with Ultracomp ebeads. The stained cells were analyzed using the FACSAria Fusion Flow cytometer (BD Biosciences).

#### RNA extraction and reverse transcription-quantitative polymerase chain reaction (RT-qPCR) from blood and immune cells

Footpads was collected in TRIzol reagent (ThermoFisher) and RNA was extracted according to manufacturer’s protocol and store at -80 C until use. RT-qPCR was performed using the NEB Luna Universal One-Step RT-qPCR kit with SYBR-Green reagent (NEB, MA, USA). Primers were ordered from Integrated DNA Technologies (IDT, Iowa, USA) and their sequence information is presented in Table S1. The reaction conditions were 55°C for 10 min for reverse transcription, 95°C for 1 min for initial denaturation and polymerase activation, 95°C for 10 seconds for denaturation, and 60°C for 30 seconds for annealing/extension by 45 cycles. Relative gene expression was determined by normalizing with GAPDH gene followed by the two delta-delta Ct (2^Ct) method of relative quantification.

#### Single cell suspension preparation for single cell RNA sequencing

We fixed CD45+ cells using Evercode cell fixation V2 kit (ECF2001, Parse Biosciences) by following standard fixation protocol provided and detailed step by step protocol has been published previously ^79^. The fixed cells were preserved at -80 °C. Then we sent samples on dry ice from Virginia Tech to the University of Michigan for library preparation and sequencing.

#### Library preparation, single cell RNA sequencing and data analysis

We used Evercode’s whole transcriptome kit (EC-W01030, Parse Biosciences) for library preparation and single cell RNA sequencing according to the manufacturer’s instructions. Briefly, we loaded 8333 fixed cells from each blood and footpad samples for three rounds of barcoding followed by lysis to isolate barcoded cDNA and prepare sub libraries. These sub libraries were sequenced on an Illumina Novaseq sequencer and generated 30-60 thousand reads per cell. The sequenced data was processed by using Parse Biosciences processing pipeline (v0.9.6p). We aligned sequencing reads to the GRCm38 mouse genome with default settings to demultiplex samples. Briefly, each of the twelve sub libraries was processed individually using the command split-pipe -mode all, and the output was combined by using split-pipe -mode combine.

Downstream processing of output count, gene/feature, and barcode matrices was performed with R/BioConductor package Seurat (v4.0.2) at default settings unless otherwise noted. Quality control analysis was done to filter for high quality cells and cells with less than 150 or more than 7,500 detected unique genes, > 40,000 unique molecular identifiers, and >15% mitochondrial reads were excluded from analysis (all thresholds were set using empirical distributions). Doublet discrimination was performed on filtered data using DoubletFinder (v 2.0.3) and doublets were excluded from downstream analysis. Filtered datasets were divided into Seurat objects corresponding to each sample (lean mock, lean MAYV-infected, obese mock, and obese MAYV-infected) and count normalization and variance stabilization was done for each sample using the SCTransform function. Integrated data analysis was carried out using the most highly variable shared genes to find analogous populations and enable direct comparison of differential gene expression when cell cohorts. Cells were clustered with Seurat using the top 15 principal components (PCs) as determined by the elbow method with a cluster resolution of 0.8 for separate cell cohort analysis and 0.4 resolution for integrated analysis. For visualizing clusters, the Uniform Manifold Approximation and Projection (UMAP) method was used. Cluster markers were identified with FindAllMarkers function (log-transformed fold-change threshold of 0.5 for separate cohort clusters and 0.25 for integrated clusters). Wilcoxon rank-sum tests were performed to determine DEGs between lean and obese mock and MAYV-infected integrated clusters using the FindMarkers function. DEGs with p<0.05 were considered statistically significant.

#### Single Cell cluster annotation

Single-cell clusters were annotated based on cluster marker gene enrichment with predefined immune cell and pathway gene sets as previously described^80^. Curated gene sets were generated from information available in literature, Mouse Genome Informatics (MGI) gene ontology (GO) terms, and immune cell-specific expression collected from the Immunological Genome Project Consortium (ImmGen). Two-sided Fisher’s Exact Test in R [fisher.test()] was used to determine the statistical differences in cluster marker gene enrichment with curated gene sets. Enrichments with p<0.05 were considered statistically significant.

#### Footpad macrophage sub clustering

Lean and obese mice footpad macrophages were selected for sub clustering. Five clusters from lean and eight clusters form obese mice expressing CD11b cell markers were characterized further using Seurat’s subset function as described above and reanalyzed similarly to the main dataset, including running the RunPCA, FindAllMarkers function (resolution = 0.5), FindMarkers function, FindClusters, and RunUMAP functions. Ambiguous cells from the subset were removed, and annotations for the remaining clusters were added to the main dataset. Each macrophage subset composition in lean and obese mice was calculated through dividing the number of specific subset cells by the total percentages of each cell subset either in lean or obese host. For each macrophage subset, differential gene expression analysis was performed on the lean infected cells from the lean mock infected, and obese infected cells from the obese mock infected cells. For summary analyses, clusters were grouped as follows: lean hosts; MF.103-11B+SALM3, MF.103-11B+, MF.103CLOSER, MF.F480HI.GATA6KO, MFIO5.II+480LO, and obese host; MF.103-11B+, MF.103-11B+24-, MF.103CLOSER, MF.F480HI.CTRL, MF.F480HI.GATA6KO, MF.RP, MFIO5.II-480HI, MFIO5.II+480INT.

#### Statistical analysis

All statistics were performed using the GraphPad version 9 and data are presented as mean ± standard deviation. The statistical tests used to analyze data are described in figure legends.

