## Supplementary figure 1 for "Obesity-Associated Changes in Immune Cell Dynamics During Alphavirus Infection Revealed by Single Cell Transcriptomic Analysis"

**Supplementary figure 1. Cytokine expression in before and after MAYV infection in lean and obese mice.** Mice were bled prior to or 2 days post-MAYV infection and cytokines were measured by multiplex Luminex assay. Comparisons were made using unpaired t tests; *p<0.05, **p<0.01, ***p<0.001, ****p<0.0001.

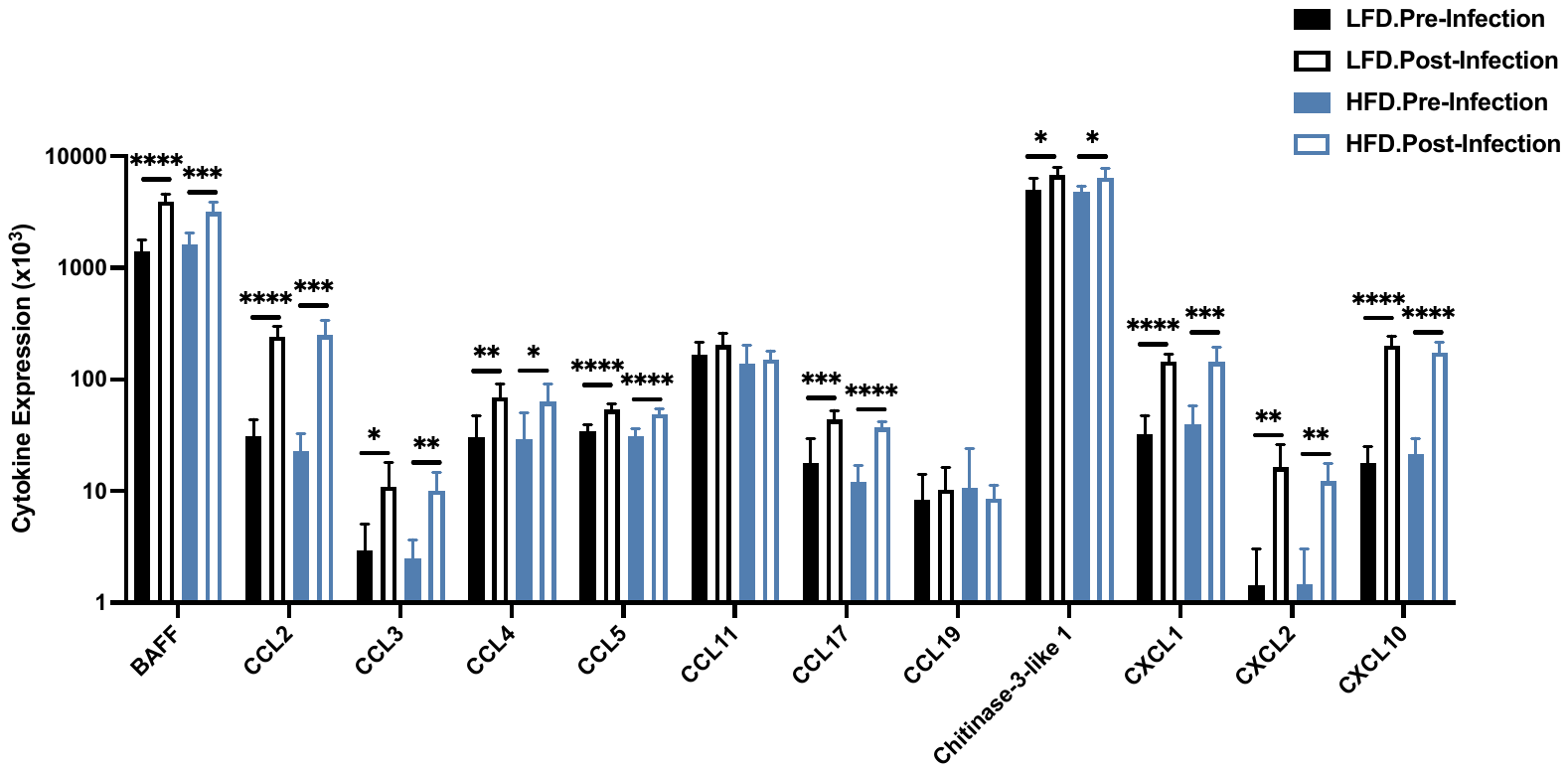

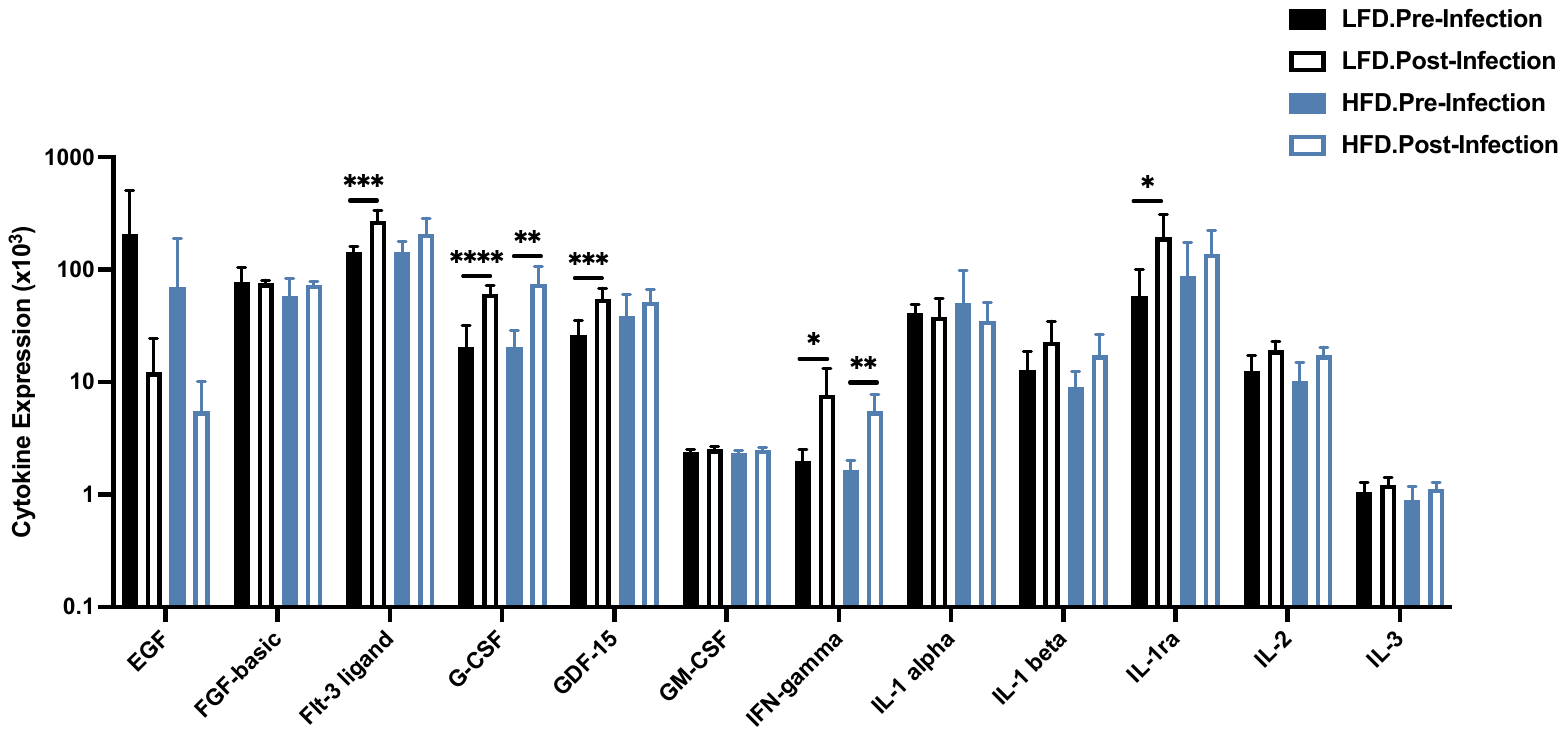

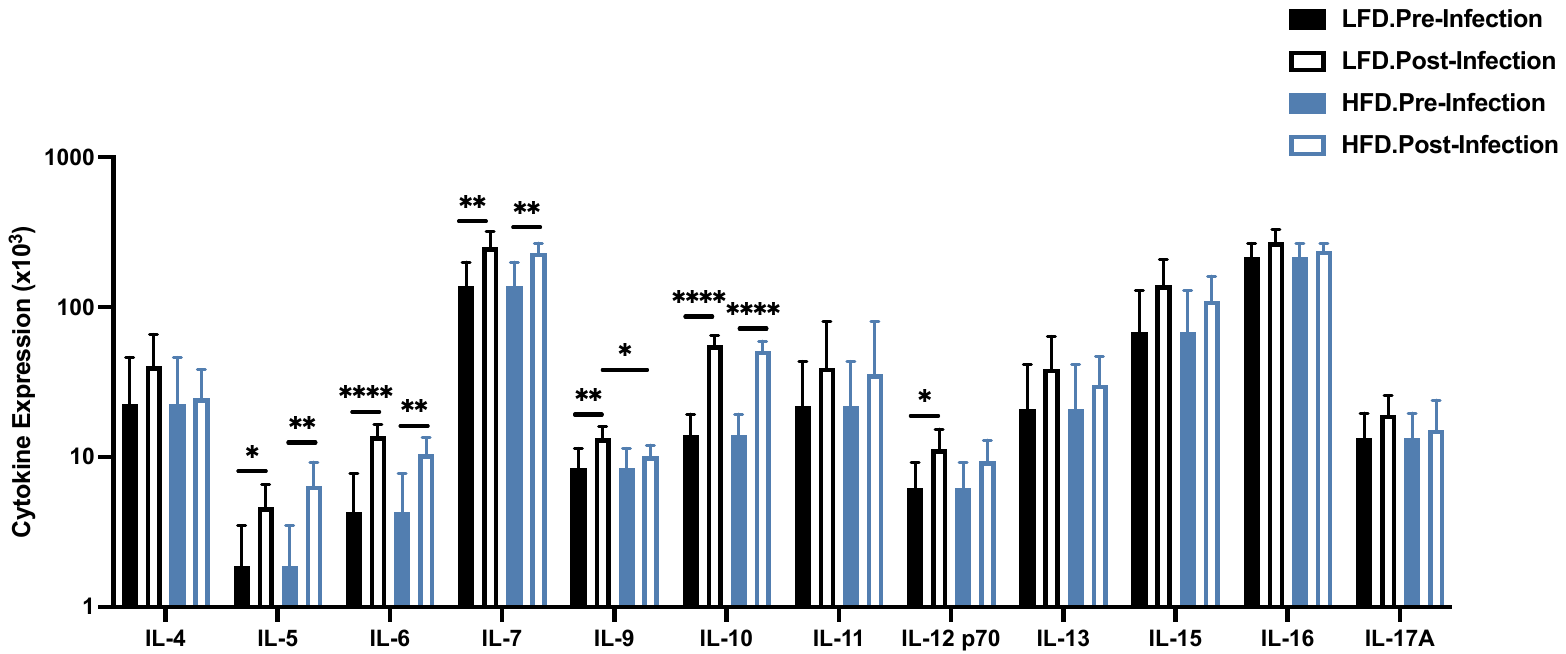

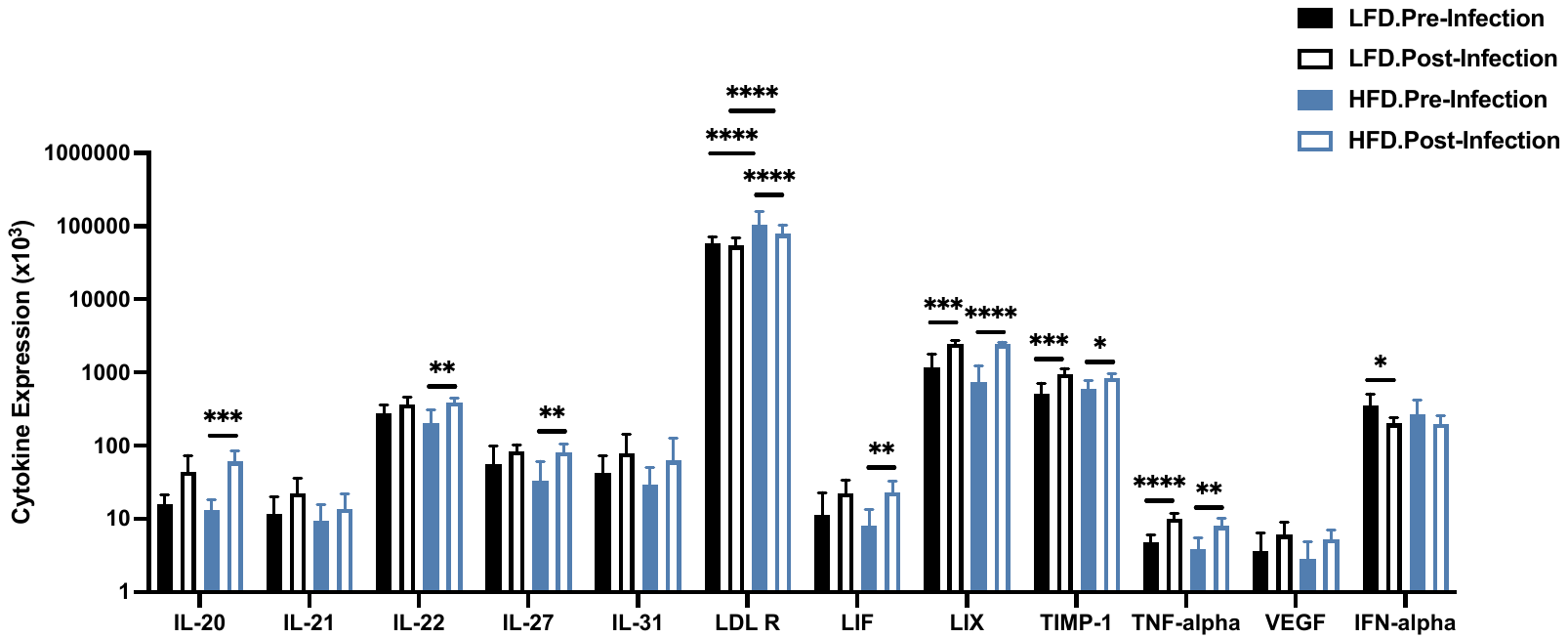
