## Supplementary figure 2 for "Obesity-Associated Changes in Immune Cell Dynamics During Alphavirus Infection Revealed by Single Cell Transcriptomic Analysis"

**Supplementary figure 2. Antiviral gene expression in lean and obese MAYV infected mice footpads.**

C57BL/6J lean and obese mice were infected with 10^4^ PFU of MAYV strain TRVL 4675 in each hind footpad and footpads were collected at 7 days post-inoculation. RNA was extracted and subjected to RT-qPCR. Data were normalized to GAPDH and analyzed using unpaired t-tests with Welch’s correction. The error bars represent the standard deviation and bars indicate mean values.

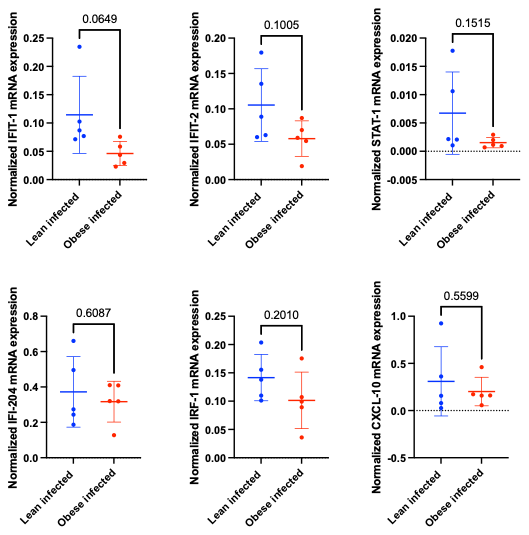
