## Supplementary figure 3 for "Obesity-Associated Changes in Immune Cell Dynamics During Alphavirus Infection Revealed by Single Cell Transcriptomic Analysis"

**Supplementary figure 3. Blood immune cell clusters detected at 2 dpi from mock- and MAYV-infected lean and obese mice.**

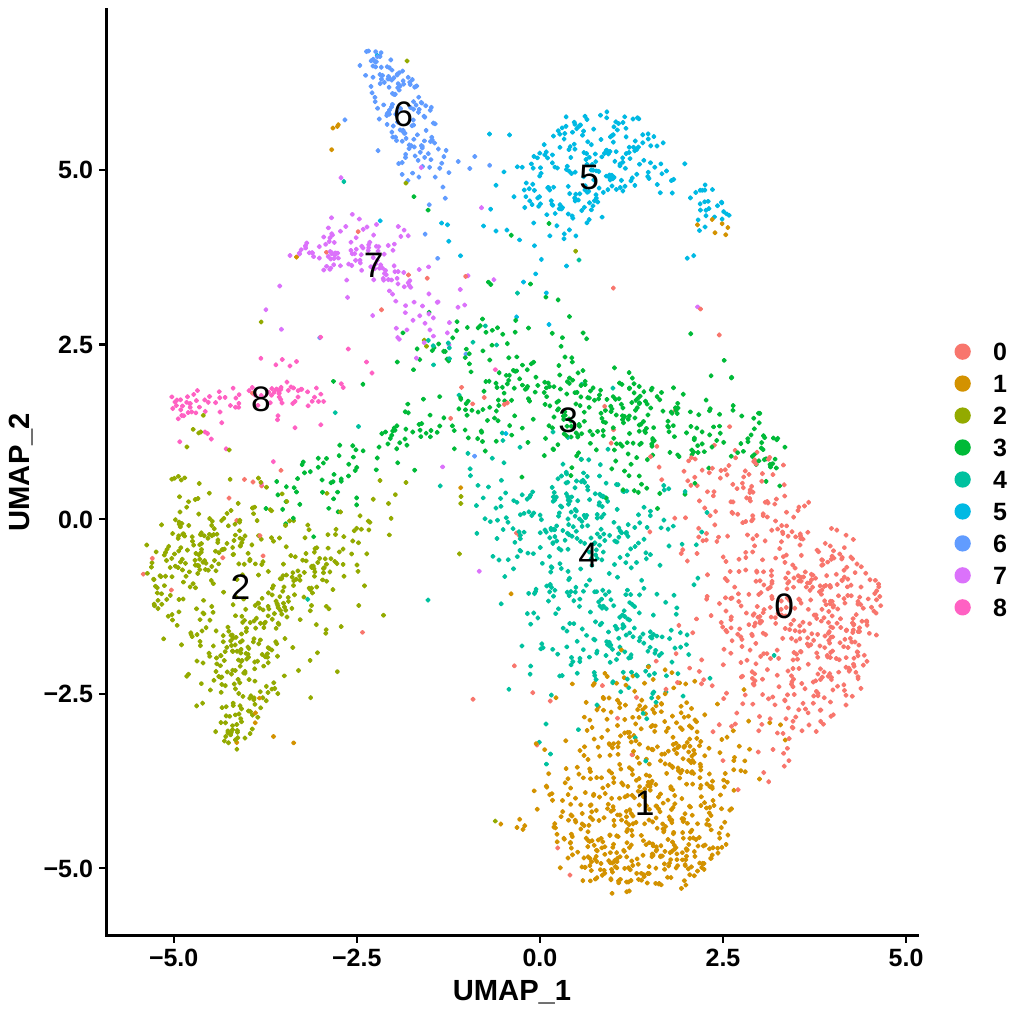

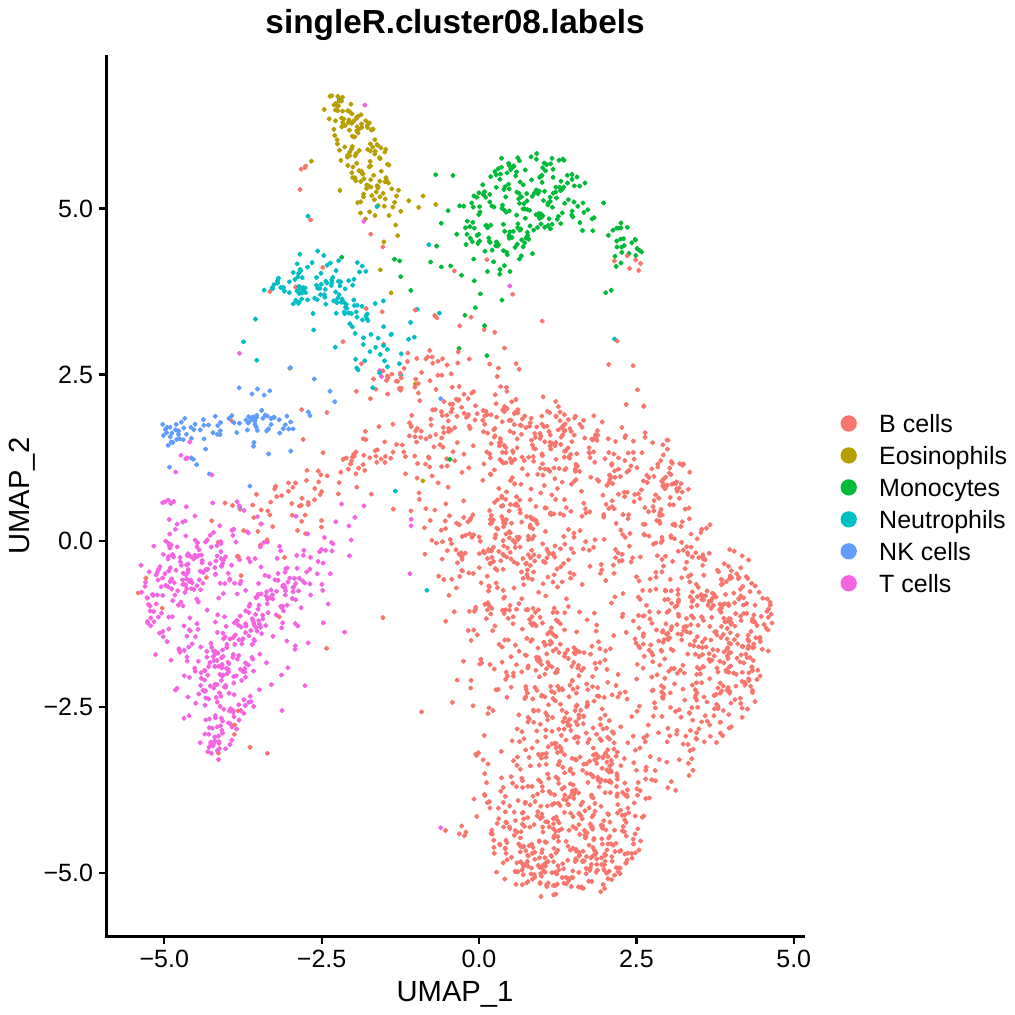
**Immune cell clusters in lean mock-infected mice (3292 cells)**

**Immune cell clusters in lean MAYV-infected mice (3640 cells)**

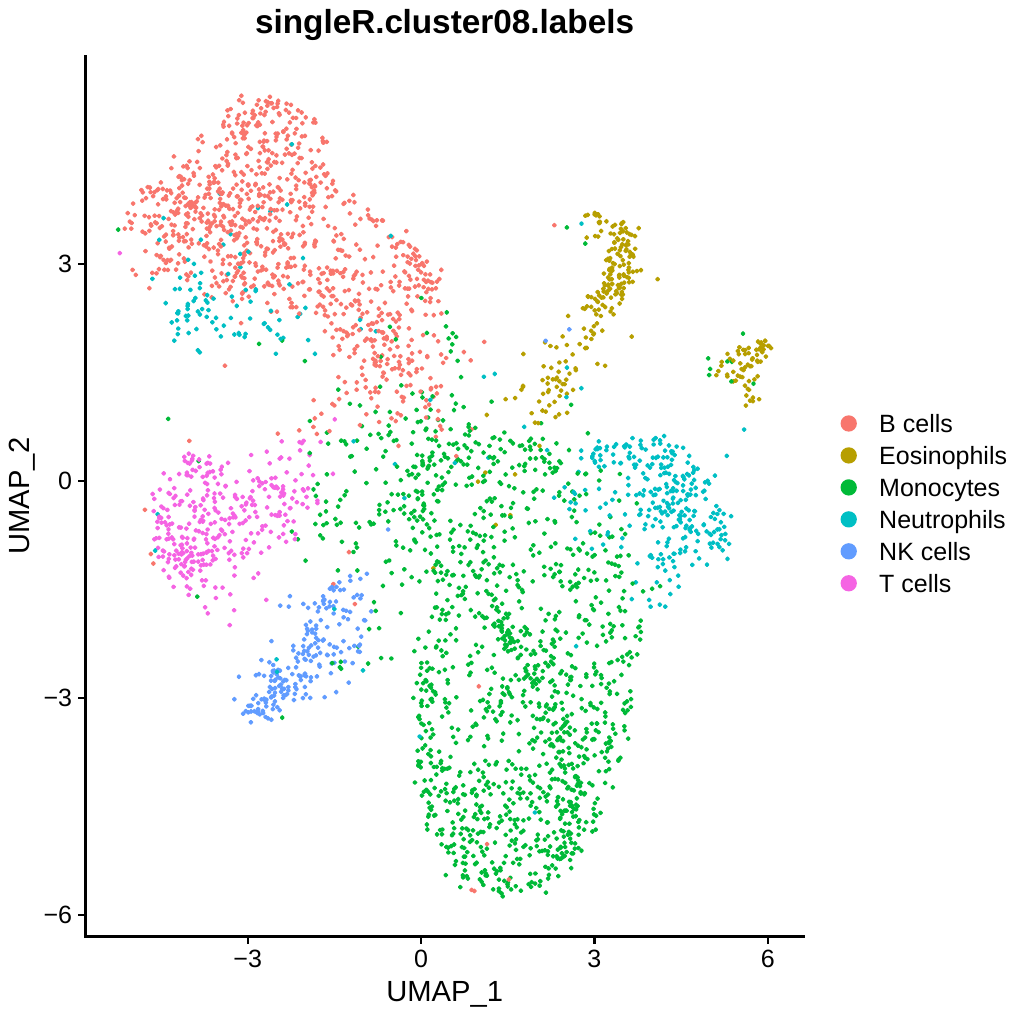

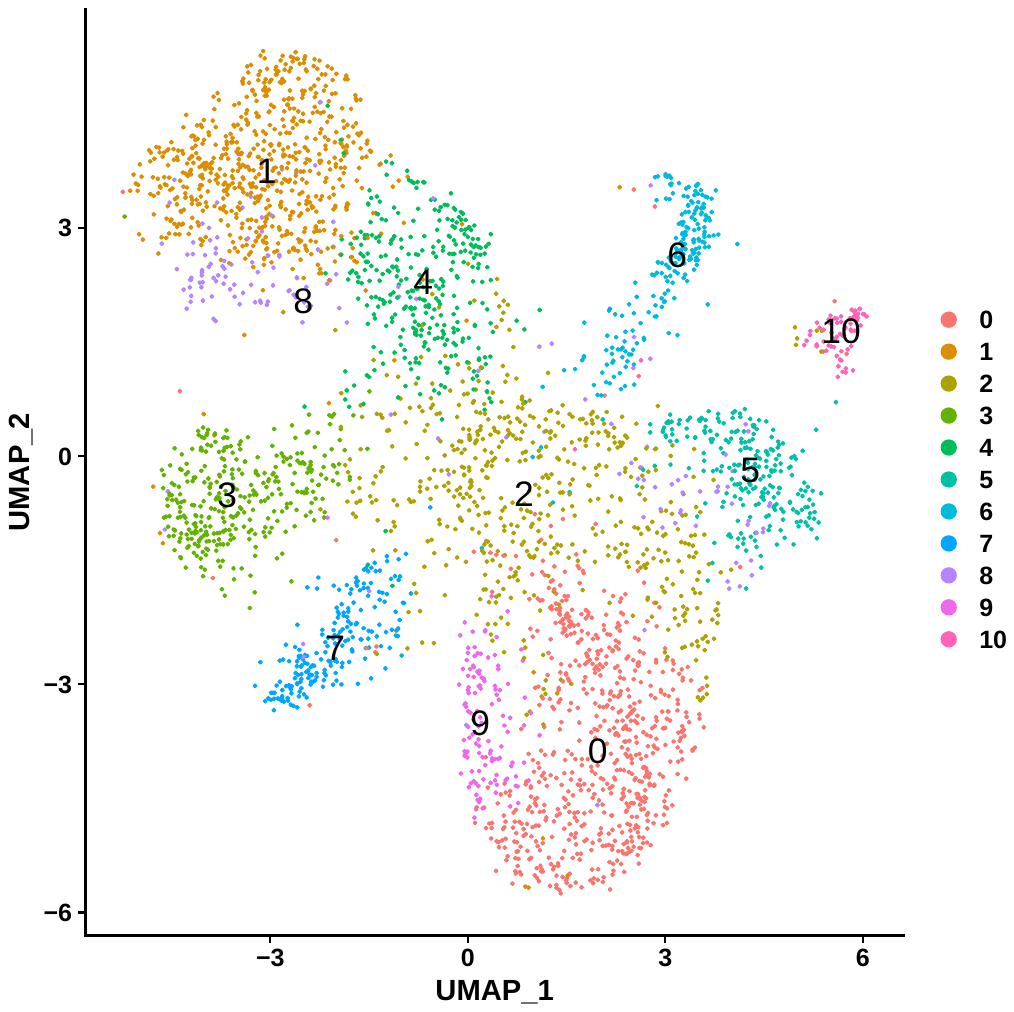

**Immune cell clusters in obese mock-infected mice (5398 cells)**

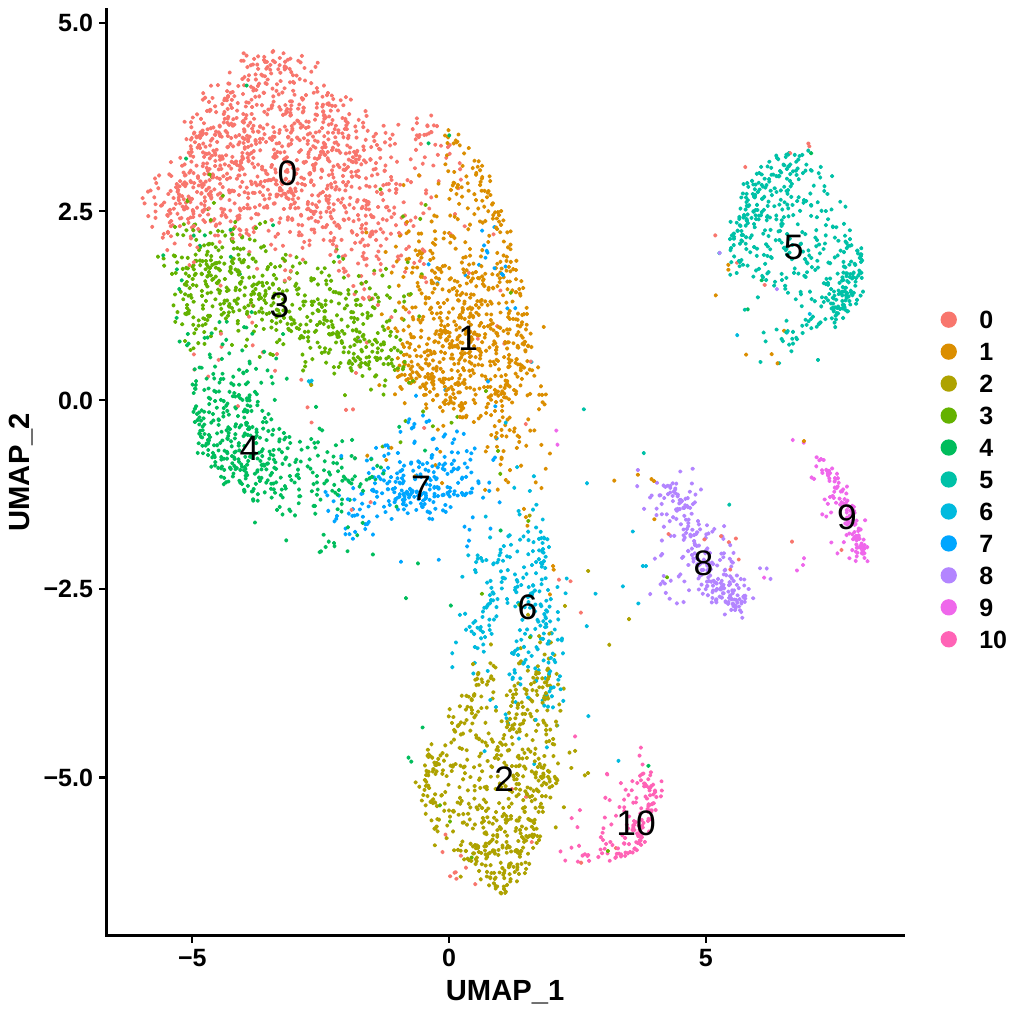

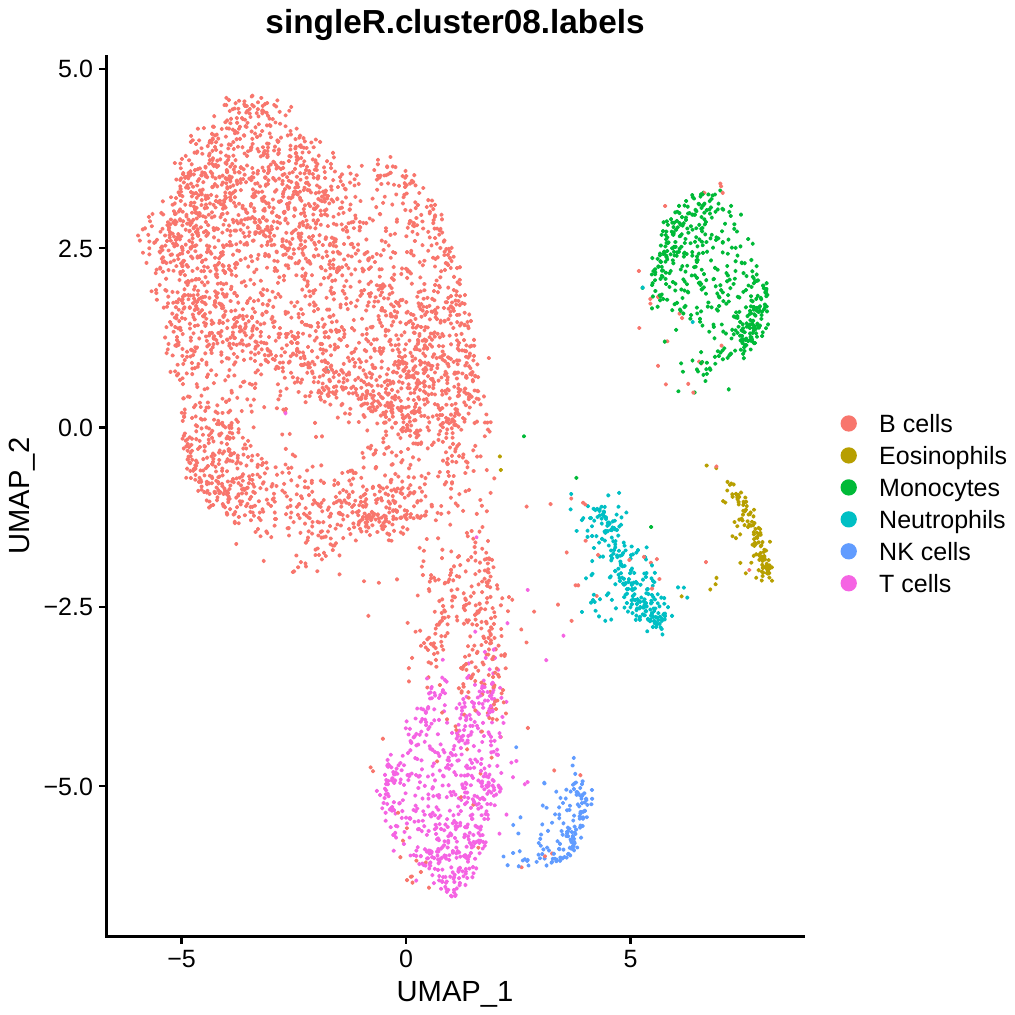

**Immune cell clusters in obese MAYV-infected mice (5960 cells)**

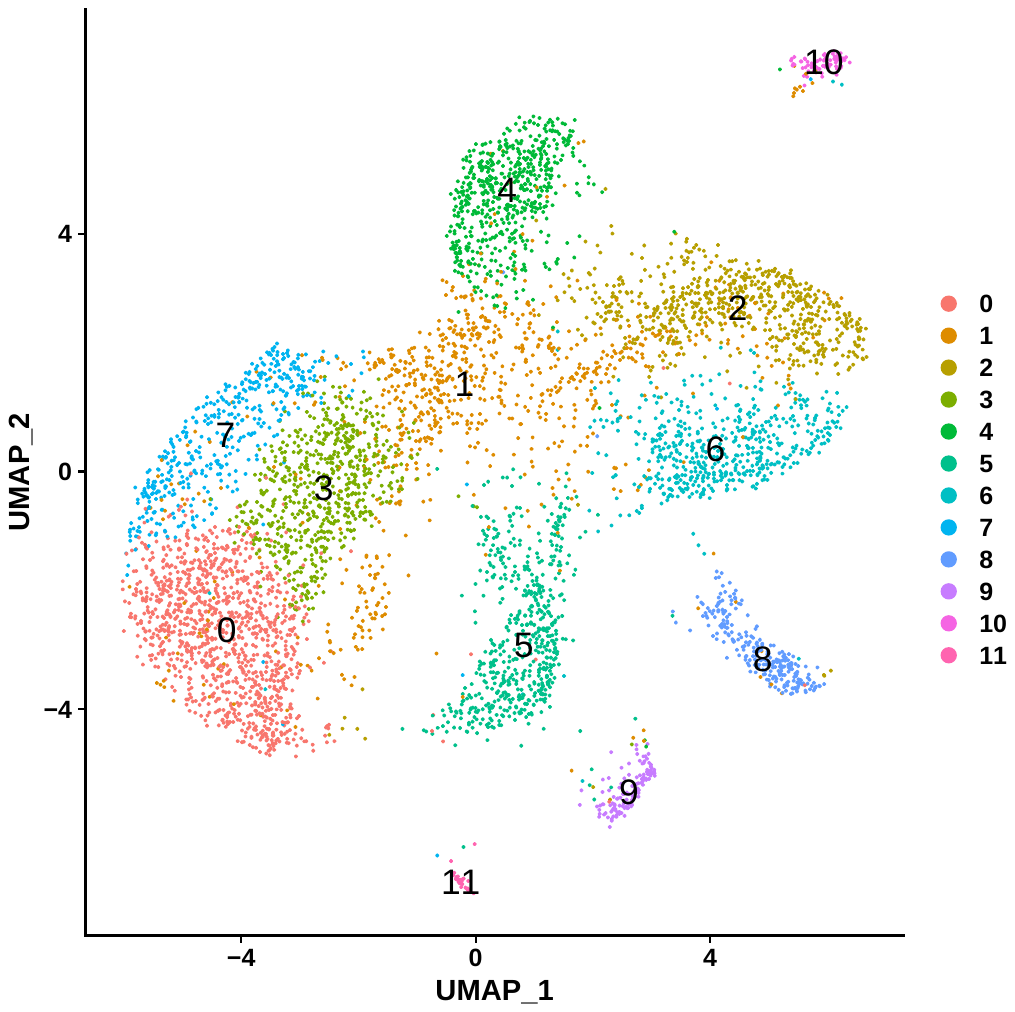

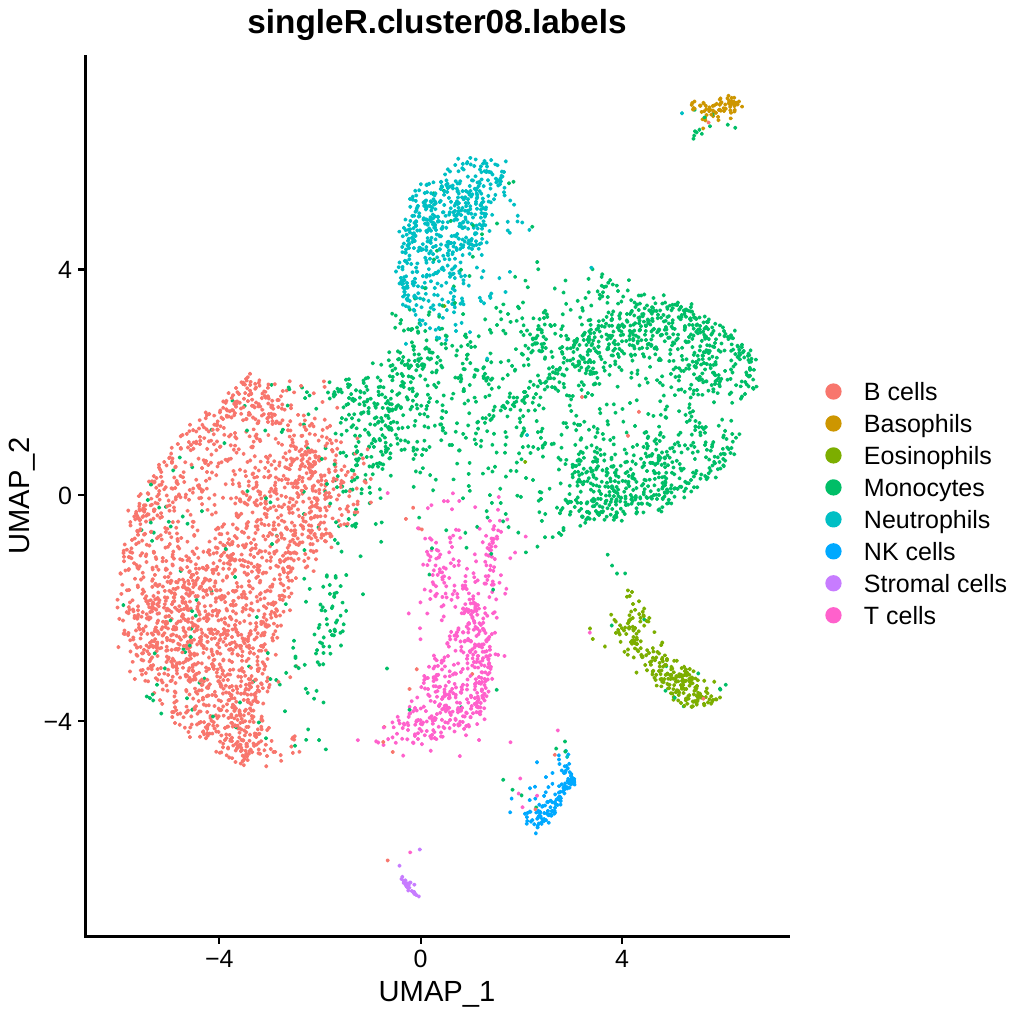
