## Supplementary figure 4 for "Obesity-Associated Changes in Immune Cell Dynamics During Alphavirus Infection Revealed by Single Cell Transcriptomic Analysis"

**Blood immune cell clusters detected at 7 dpi from mock- and MAYV-infected lean and obese mice.**

**Immune cell clusters in lean mock infected mice blood at 7 dpi (Total cells 5087)**

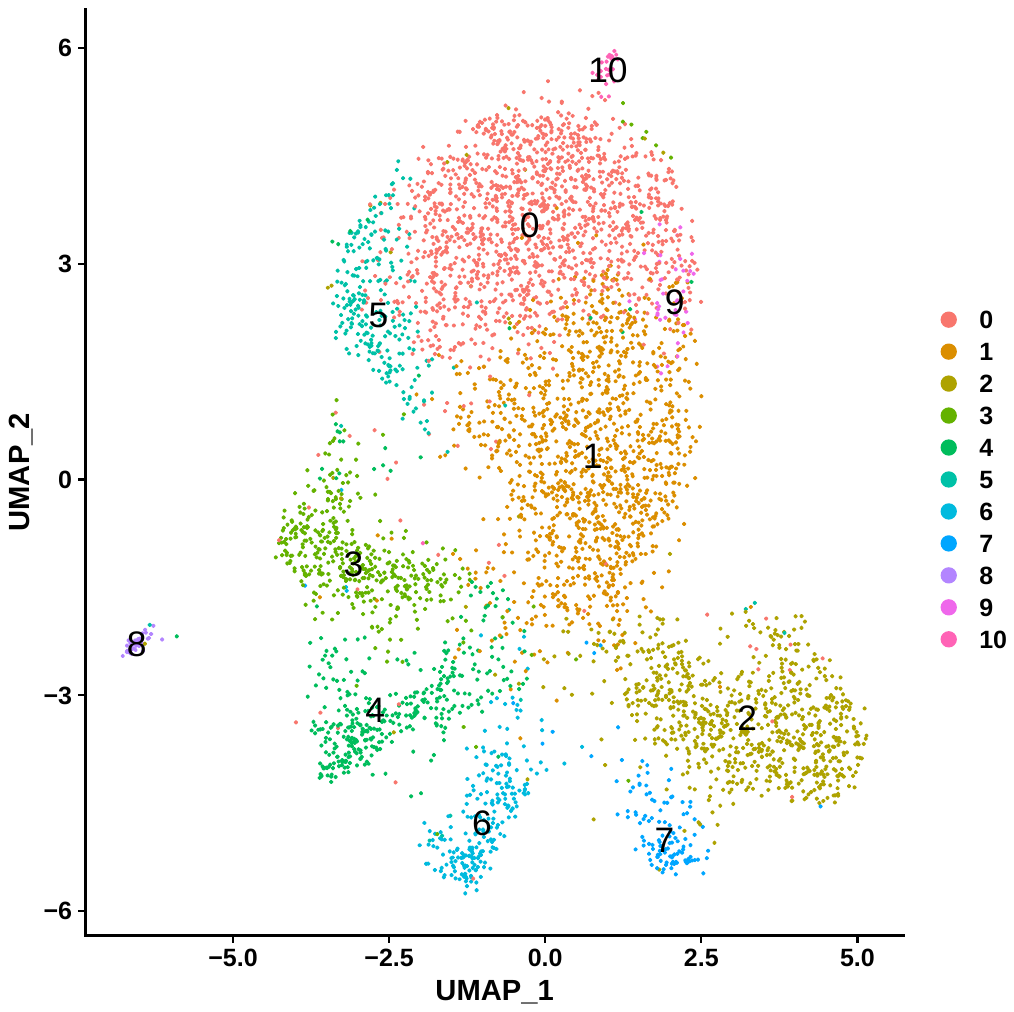

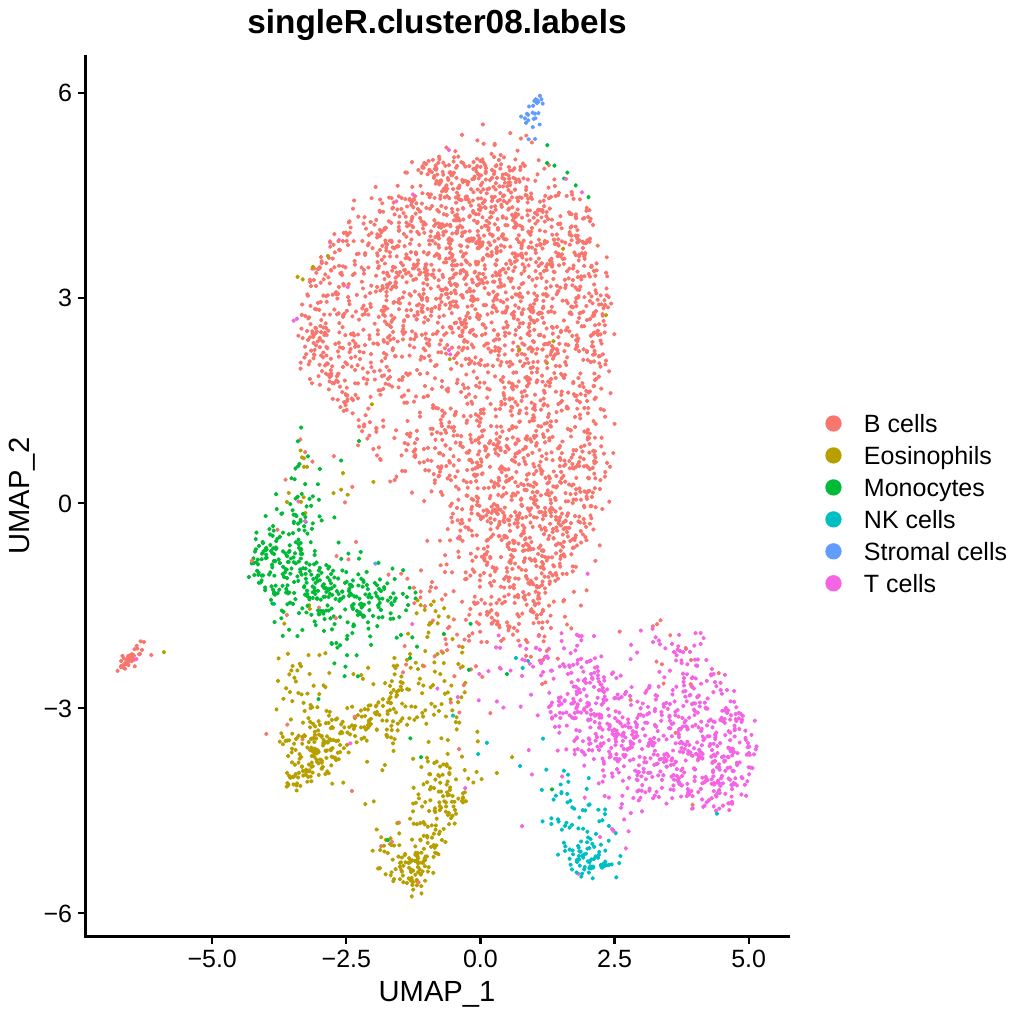

**Immune cell clusters in lean MAYV-infected mice blood at 7 dpi (Total cells 5307)**

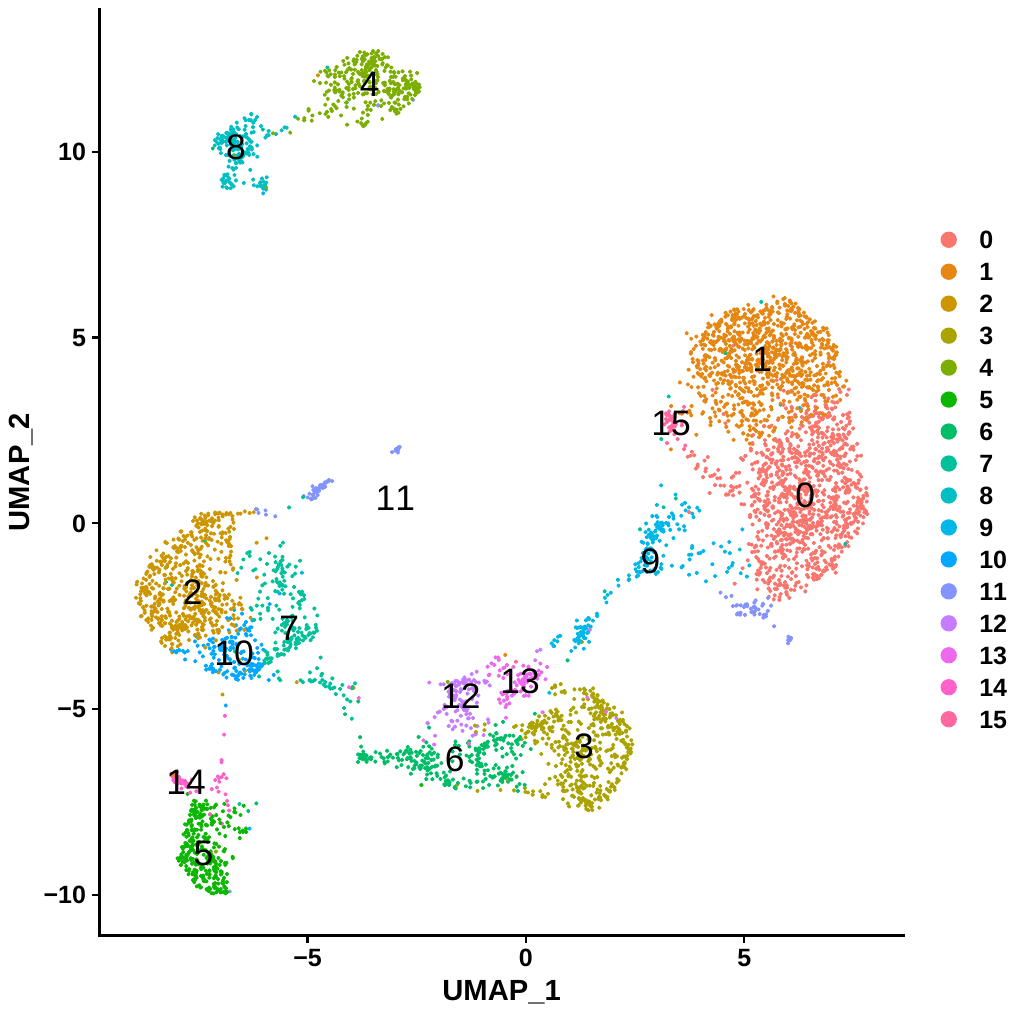

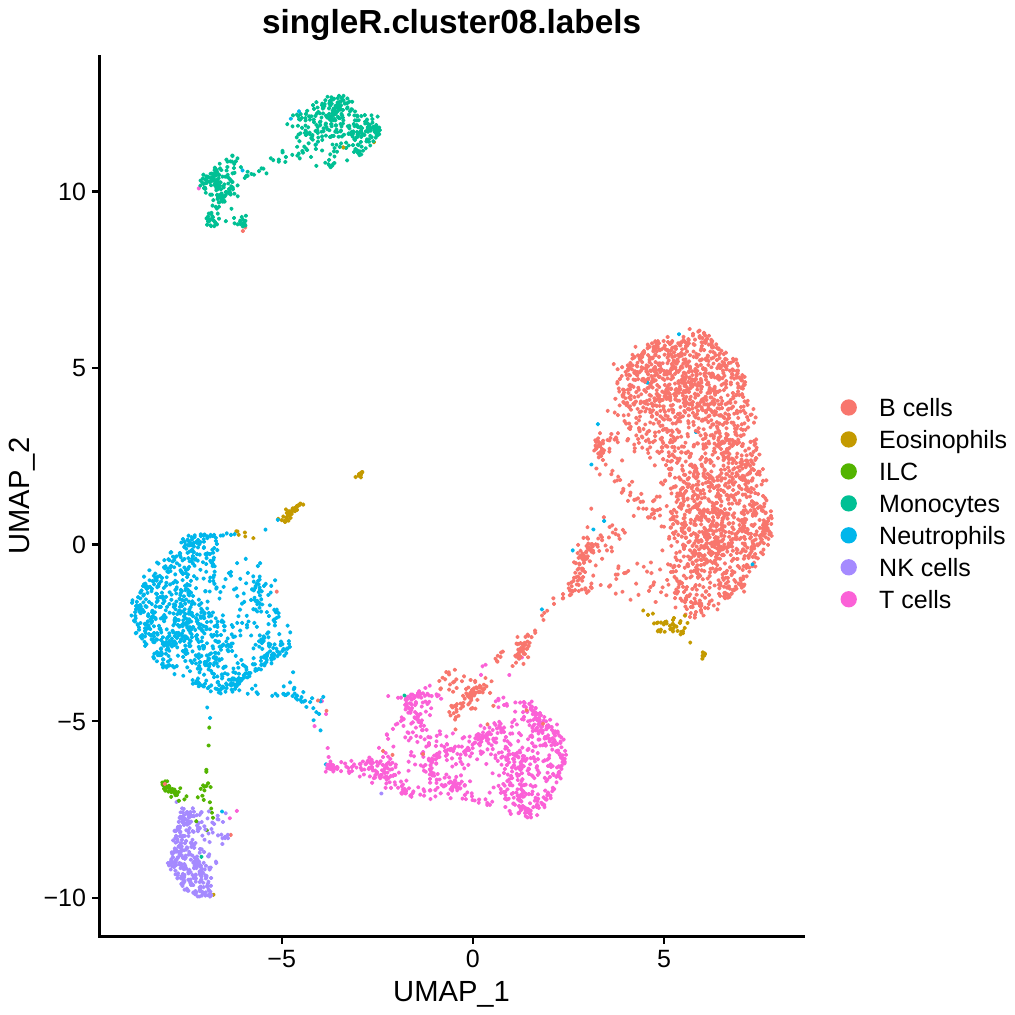

**Immune cell clusters in obese mock infected mice blood at 7 dpi (Total cells 6113)**

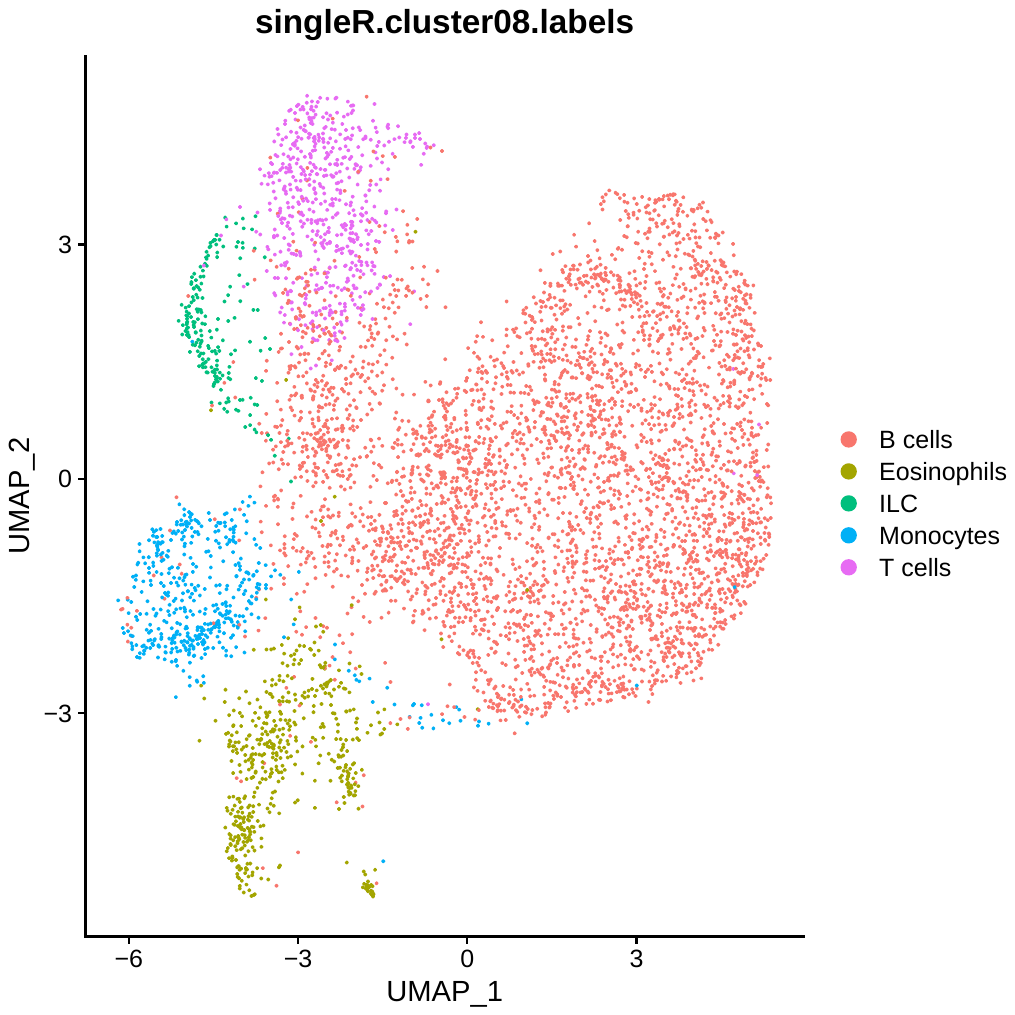

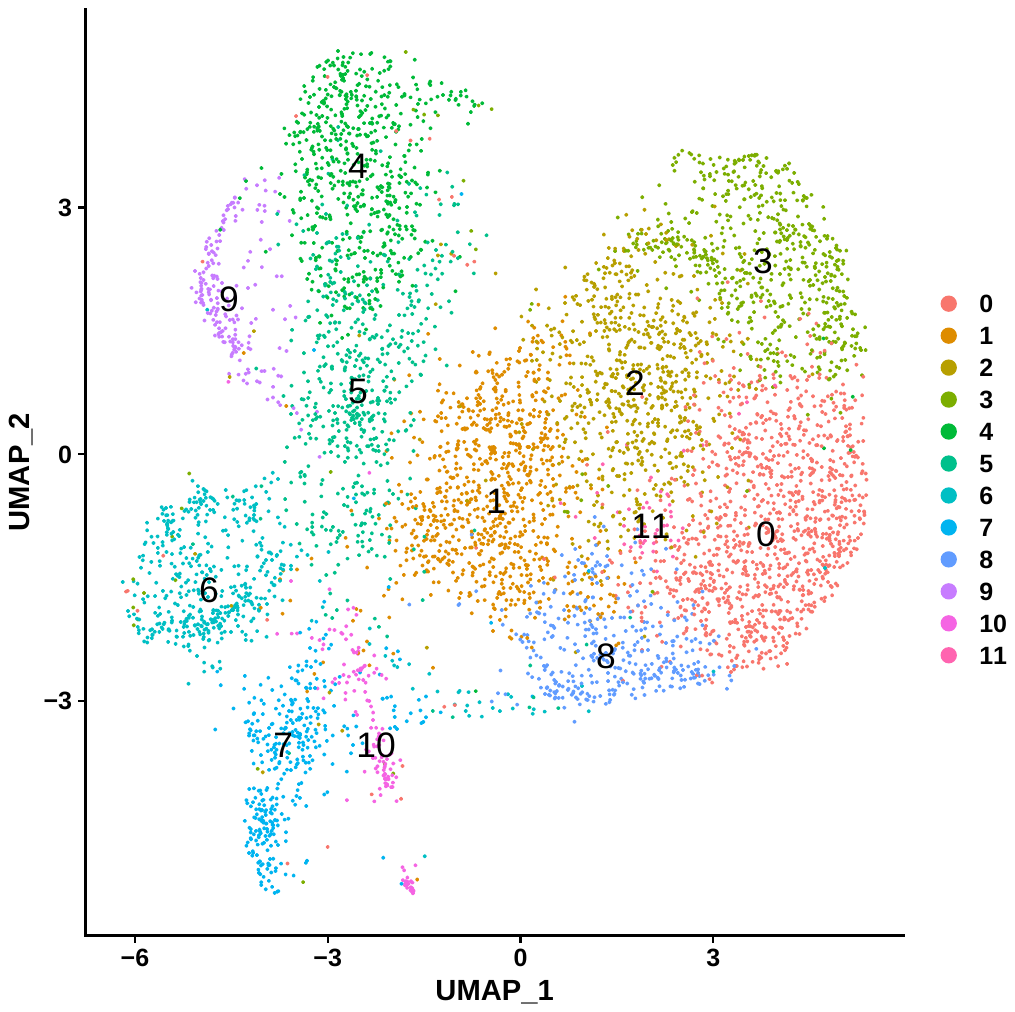

**Immune cell clusters in obese MAYV-infected mice blood at 7 dpi (Total cells 5606)**

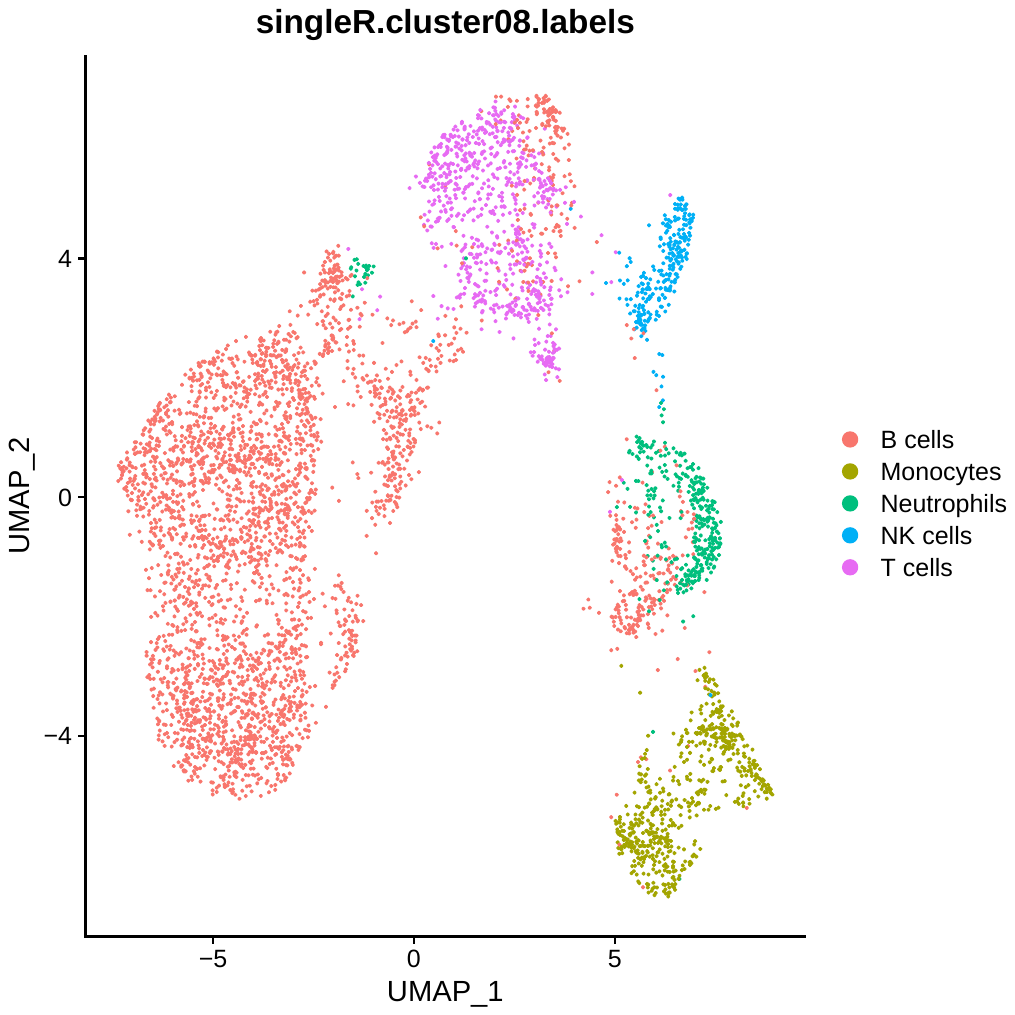

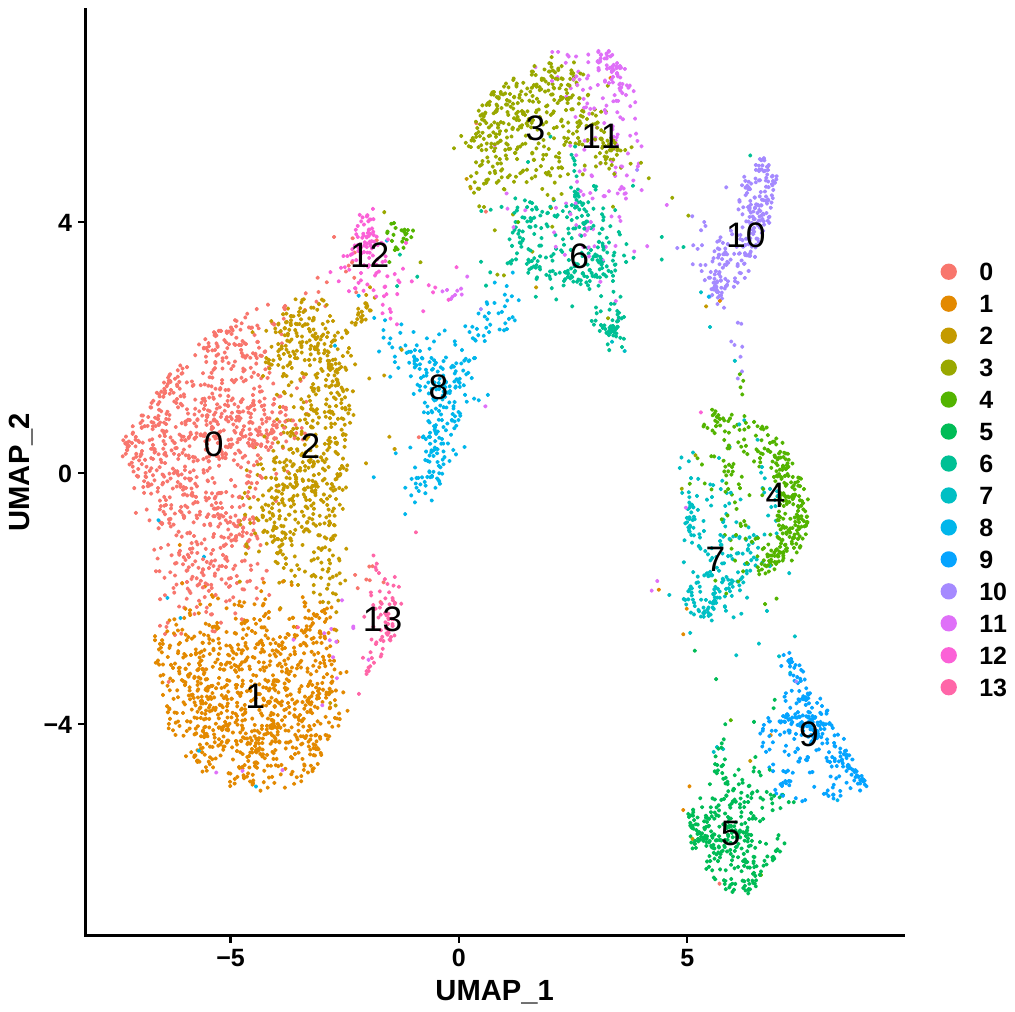
