## Supplementary Table 1 for "Obesity-Associated Changes in Immune Cell Dynamics During Alphavirus Infection Revealed by Single Cell Transcriptomic Analysis"

**Supplementary table 1. List of RT-qPCR primers used to assess gene expression.**

| **Primer target** | **Sense** | **Sequence 5' → 3'** |
| --- | --- | --- |
| IFN-α | F | GGATGTGACCTTCCTCAGACTC |
|  | R | ACCTTCTCCTGCGGGAATCCAA |
| GAPDH | F | AGGTCGGTGTGAACGGATTTG |
|  | R | TGTAGACCATGTAGTTGAGGTCA |
| IFN-β, | F | CAGCTCCAAGAAAGGACGAAC |
|  | R | GGCAGTGTAACTCTTCTGCAT |
| IFN-γ | F | ATGAACGCTACACACTGCATCTTG |
|  | R | GCAGCGACTCCTTTTCCGC |
| Rsad-2 | F | GGAAGGTTTTCCAGTGCCTCCT |
|  | R | ACAGGACACCTCTTTGTGACGC |
| CXCL-10 | F | ATGAACCCAAGTGCTGCCGT |
|  | R | AGGAGCCCTTTTAGACCTTTTTTG |
| GBP2 | F | CTGCACTATGTGACGGAGCTA |
|  | R | CGGAATCGTCTACCCCACTC |
| IFIT-1 | F | TACAGGCTGGAGTGTGCTGAGA |
|  | R | CTCCACTTTCAGAGCCTTCGCA |
| IFIT-2 | F | CGAACTACCGTCTGGATGACTG |
|  | R | CTTCAACCAGCGCCATTGCTTG |
| IFIT-3b | F | GCTCAGGCTTACGTTGACAAGG |
|  | R | CTTTAGGCGTGTCCATCCTTCC |
| IL-6 | F | TAGTCCTTCCTACCCCAATTTCC |
|  | R | TTGGTCCTTAGCCACTCCTTC |
| MX-1 | F | TGGACATTGCTACCACAGAGGC |
|  | R | TTGCCTTCAGCACCTCTGTCCA |
| IRF-1 | F | TCCAAGTCCAGCCGAGACACTA |
|  | R | ACTGCTGTGGTCATCAGGTAGG |
| STAT-1 | F | GCCTCTCATTGTCACCGAAGAAC |
|  | R | TGGCTGACGTTGGAGATCACCA |

**F stands for forward primer; R stands for reverse primer.
